# Monoclonal anti-AMP-antibodies reveal broad and diverse AMPylation patterns in cancer cells

**DOI:** 10.1101/2020.06.23.164731

**Authors:** Dorothea Höpfner, Joel Fauser, Marietta S. Kaspers, Christian Pett, Christian Hedberg, Aymelt Itzen

**Affiliations:** Department of Biochemistry and Signaltransduction, University Medical Center Hamburg-Eppendorf (UKE), Martinistr. 52, 20246, Hamburg, Germany; Center for Integrated Protein Science Munich (CIPSM), Department Chemistry, Technical University of Munich, Lichtenbergstrasse 4, 85747, Garching, Germany; Chemical Biology Center (KBC), Department of Chemistry, Umeå University, Linnaeus väg 10, 90187, Umeå, Sweden; Center for Structural Systems Biology (CSSB), University Medical Center Hamburg-Eppendorf (UKE), Hamburg, Germany

## Abstract

AMPylation is a post-translational modification that modifies amino acid side chains with adenosine monophosphate (AMP). Recent progress in the field reveals an emerging role of AMPylation as a universal regulatory mechanism in infection and cellular homeostasis, however, generic tools to study AMPylation are required. Here, we describe three monoclonal anti-AMP antibodies (mAbs) from mouse which are capable of protein backbone independent recognition of AMPylation, in denatured (Western Blot) as well as native (ELISA, IP) applications, thereby outperforming previously reported tools. These antibodies are highly sensitive and specific for AMP modifications, highlighting their potential as tools for new target identification, as well as for validation of known targets. Interestingly, applying the anti-AMP mAbs to various cancer cell lines reveals a previously undescribed broad and diverse AMPylation pattern. In conclusion, the anti-AMP mABs will aid the advancement of understanding AMPylation and the spectrum of modified targets.

## Introduction

Post-translational modifications (PTMs) are diverse covalent alterations that modulate the activity, localization, stability, and specificity of proteins. One such PTM is AMPylation (also referred to as adenylylation), occurring in prokaryotes as well as eukaryotes. Enzymes utilize adenosine triphosphate (ATP) as donor substrate to transfer the adenosine monophosphate (AMP) to hydroxyl-bearing amino acid side chains (e.g. tyrosine, serine, threonine) of a target protein, with pyrophosphate being released as a side product. There are three different classes of AMPylators or protein adenylyl transferases known to date: DNA-β-Polymerase-like AMPylators with their most prominent member DrrA from *Legionella pneumophila*, FIC (*Filamentation induced by cyclic AMP*) enzymes represented by human HYPE/FICD (Engel et al., 2012), IbpA from *Histophilus somni* (Worby et al., 2009a), or VopS from *Vibrio parahaemolyticus* (Yarbrough et al., 2009), and - as most recent discovery - pseudokinases, specifically the highly conserved SelO (Sreelatha et al., 2018).

AMPylation has been studied over 50 years (Kingdon et al., 1967), and has gained recent attention with the identification of small GTPases as targets of AMPylating enzymes during various bacterial infections (Worby et al., 2009a; Yarbrough et al., 2009). The discovery of FICD/HYPE as the only mammalian FIC protein and its modification of the endoplasmic reticulum (ER) chaperone Bip (Engel et al., 2012; Ham et al., 2014) illustrates a role of AMPylation in protein homeostasis (Preissler et al., 2015, 2016; Sanyal et al., 2015). Recent findings on AMPylation by pseudokinases (Sreelatha et al., 2018) hints at a broader occurrence of this modification as a general mechanism, and not just in context of bacterial infections as previously thought.

However, despite a high prevalence of predicted FIC enzymes based on their conserved sequence, especially in pathogenic bacteria (Khater & Mohanty, 2015), only a limited amount of AMPylation targets is known. This discrepancy between number of enzymes and identified targets highlights the challenge of detecting AMPylation. Available tools are limited and associated with disadvantages when it comes to necessary resources and/or studying AMPylation in a physiologically relevant context. ATP analogs have reduced intracellular uptake (Plagemann & Wohlhueter, 1980) (although recent work established a cell permeable pronucleotide probe (Kielkowski et al., 2020)), are competed by the high endogenous amounts of ATP, and hampered by the potential inability of enzymes to use these analogs as substrates. When used in cell lysates, spatial and temporal regulation is abrogated.

Antibodies targeting AMPylation could overcome many of these challenges as well as offer further applications as an orthogonal approach. Ideally, such an antibody would be able to detect AMPylation with high sensitivity and specificity in native as well as denatured proteins, thus enabling Western Blot (WB) detection as well as enrichment by IP. Previously generated polyclonal antibodies using AMPylated peptides do not fulfill the desired properties (Hao et al., 2011; Smit et al., 2011).

Here, we generate three new monoclonal antibodies from mice, recognizing AMPylation independently of the protein backbone, under denatured as well as native conditions. Besides validation of targets, they can serve as new tools for target identification. Since new target identification hinges on proper positive controls, and false negatives may not be detected, a thorough characterization of the antibodies’ behavior in the specific application is crucial.

## Results

Previously published and commercially available antibodies claimed to recognize AMPylated threonine and tyrosine, respectively, independent of the peptide backbone and protein environment (see Sigma-Aldrich 09-890 and ABS184)(Hao et al., 2011). However, evaluation of their performance in Western Blot on various recombinant proteins with different modified amino acid side chains [such as Rho GTPase Cdc42 AMPylated at threonine35 (VopS (Yarbrough et al., 2009)) or tyrosine32 (IbpA (Worby et al., 2009b)), respectively, Rab GTPase Rab1b modified at tyrosine77 (DrrA (Müller et al., 2010)), Histone H3.1 modified at threonine (HYPE (Truttmann et al., 2016)), the ER chaperone Bip/Grp-78 modified at threonine518 (HYPE (Preissler et al., 2015)) and the FIC enzyme HYPE/FICD auto-modified at threonine80,183+serine79 (Sanyal et al., 2015)] showed that they do not recognize all AMPylations (Figure 1A): While the commercially available anti-Thr-AMP antibody (Sigma-Aldrich 09-890) successfully recognized Cdc42-Thr-AMP and Hype-Thr-AMP, the detection of H3.1-Thr-AMP and Bip-Thr-AMP was less sensitive and in case of the latter no longer possible at 50 ng. The commercially available anti-Tyr-AMP antibody (ABS184, Merck) showed unsatisfactory performance by cross reacting with unmodified Rab1b and HYPE, respectively, as well as H3.1-Thr-AMP, in addition to exhibiting a generally weak detection signal. Since both anti-AMP antibodies did not yield broad recognition of AMPylation, we wondered whether available anti-ADP-ribosylation antibodies might also be able to detect AMPylation, since both modifications share the AMP-moiety. We therefore tested the commercially available anti-pan-ADP-ribose binding reagent (MABE1016, Merck) and – while detecting mono-ADP-ribosylated PARP3 (by autocatalysis (Vyas et al., 2014)) - found it to be unable to detect AMPylation with the exception of H3.1-Thr-AMP.

**Figure 1:**
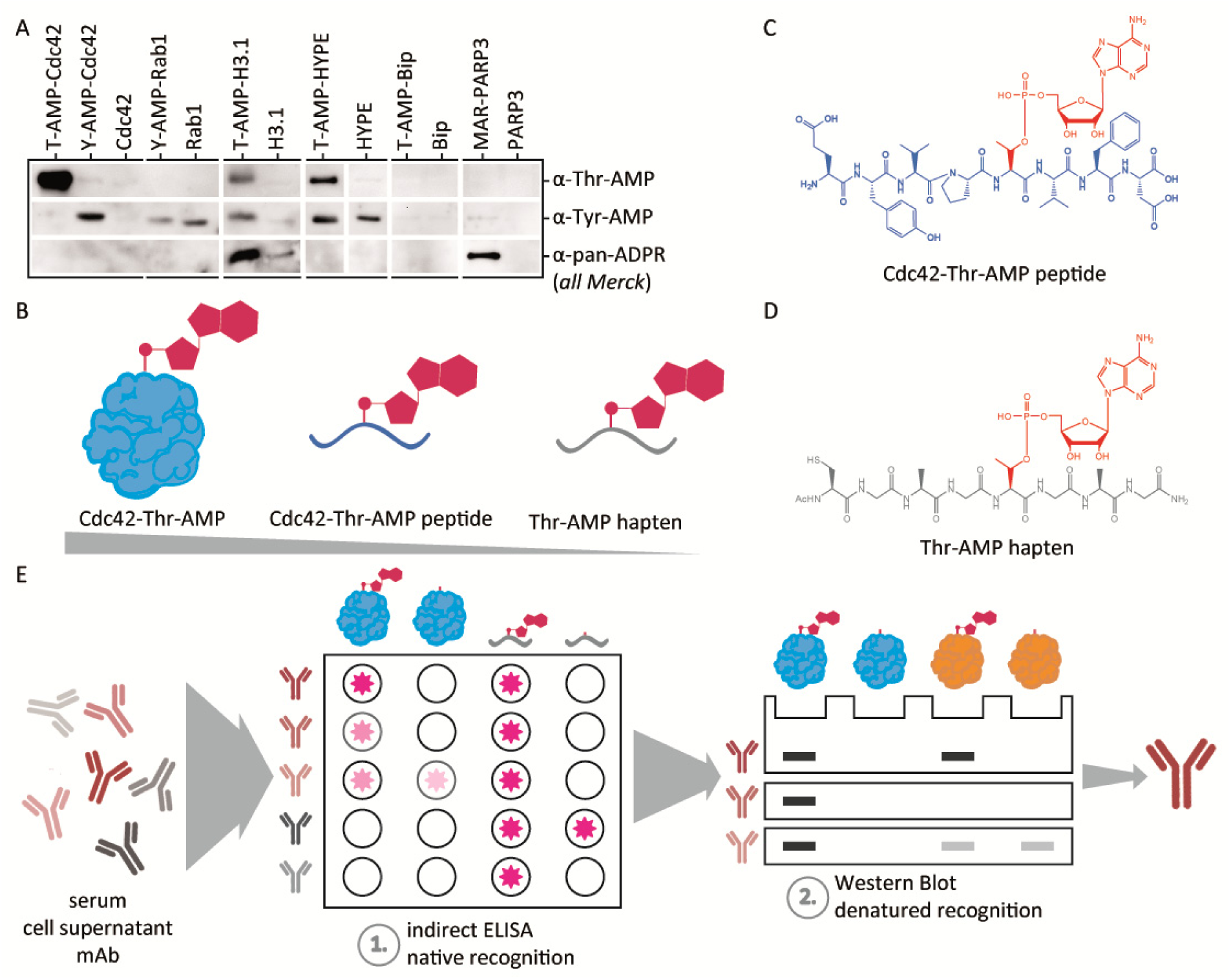
Motivation, hapten design and selection strategy for the generation of monoclonal anti-AMP antibodies. **A** Performance of commercially available AMP-antibodies. 50 ng of indicated recombinant protein was analyzed by WB using anti-threonine-AMP (Merck) and anti-tyrosine-AMP (Merck) antibodies as well as anti-ADPR binding probe (Merck) as indicated. **B** Reductive approach of hapten design. Instead of using intact AMPylated protein or AMPylated peptide from a naturally occurring target such as Cdc42, the peptide backbone’s complexity was reduced to ensure development of antibodies against the AMP-moiety alone. **C** Representation of the peptide 31-38aa in naturally occurring Cdc42-Thr-AMP with its complex side chains. **D** Representation of Thr-AMP hapten peptide with its reduced complexity of a glycine-alanine backbone, N-terminal acetylation and C-terminal amidation. A N-terminal cysteine was included to enable fusion to carrier protein. **E** Illustration of stepwise selection process of mice (sera), clones (supernatant), subclones (supernatant) and confirmation of purified antibodies during antibody generation. Candidates were first subjected to ELISA against AMPylated hapten peptide and AMPylated Cdc42-Thr, and positive clones evaluated for their performance in WB on various AMPylated proteins.

This number of false positive and of false negative signals in commercially available anti-Tyr-AMP and anti-ADPR antibodies as well as the low sensitivity in the anti-Thr-AMP antibody led us to the development of new monoclonal anti-AMP antibodies in mice.

### Design and synthesis of the AMP-bearing peptide

Previous antibodies were used and worked mostly against denatured targets in WB, and were only evaluated against small GTPases (dependent on the peptide used for immunization) (Hao et al., 2011; Smit et al., 2011). Our goal was to create a universal tool that can recognize AMPylation, not only on the rising number of known AMPylated proteins but also on unknown targets independent of backbone and protein environment, in denatured as well as native applications. This would allow for target enrichment from complex samples as well as target validation and characterization.

Instead of using an AMPylated peptide derived from a naturally occurring target protein as hapten, we chose a reductive approach, that aimed to develop the antibody against the AMP-side chain moiety alone, but not against the peptide sequence itself (Figure 1B). The strategy was therefore to reduce the peptide backbone (Figure 1C) to a non-immunogenic 8 amino acid sequence of glycine and alanine, long enough to not unintentionally cause an immune reaction to the termini, but short enough to diminish the immune response to the peptide itself. To lower the charge at the termini and simulate a natural protein peptide backbone, the peptide was N-terminally acetylated and C-terminally amidated. The AMPylated threonine was introduced in the middle of the synthesized peptide via the use of an AMPylated building block (Albers et al., 2011; Smit et al., 2011). An N-terminal cysteine was incorporated to enable fusion to carrier proteins for immunization (Figure 1D).

Since this reductive strategy of AMPylated threonine incorporated into a short glycine-alanine backbone has never been tested before and posed the risk of abolishing immunogenicity, we decided on a broad approach, choosing two different carrier proteins as hapten conjugates and two different mice breeds for immunization. In total, 10 mice were immunized with either bovine serum albumin (BSA) or keyhole limpet hemocyanin (KLH) conjugates, each of which was injected into three BALB/c and two C57 BL/6 mice by GenScript.

To ensure backbone independent recognition of AMPylation, antibody candidates were reversely selected by a stepwise screening procedure during all stages of development (Figure 1E). The screening process started with all candidates that were able to recognize the hapten with its reduced backbone complexity, proceeding to filter all candidates that were capable of recognition of native threonine-AMPylated Cdc42 as determined by ELISA (Figure 1E, 1^st^ step), and subsequently testing recognition of multiple modified proteins in denatured state via WB (Figure 1E, 2^nd^ step). Only candidates positive for all these criteria and all target proteins were taken into consideration and used for further development (Supp. Figure S1).

Our selection strategy followed by rigorous characterization aimed to overcome the aforementioned pitfalls of currently available antibodies and created three new antibodies against AMPylation: One promising clone, 17G6, with sensitive recognition of all AMPylated proteins in WB independent of their modified side chain, native recognition of Cdc42-Thr-AMP in ELISA and low background was selected for subcloning and subsequent production and purification. Another one, 7C11, was selected for showing a bias in WB for threonine modified protein. One further clone, 1G11, was selected for its development of a Tyr-AMP-specific recognition, despite immunization with a threonine-modified peptide. All monoclonal antibodies derived from C57 BL/6 mice, where 17G6’s hapten was fused to BSA while 7C11’s and 1G11’s haptens were KLH fusions (Supp. Figure S1).

### Generated anti-AMP-antibodies are highly specific for AMPylation

The three new anti-AMP antibodies 17G6, 7C11 and 1G11 were subsequently produced by roller bottle cell culture, purified from the supernatant via Protein A affinity capture, and tested for their performance in the recognition of denatured protein targets via WB. Here, sensitivity and detection limits, specificity towards AMPylation as opposed to incorporation of other nucleotides, and cross reactivity with other PTMs were evaluated. In addition, native binding as previously shown by ELISA was confirmed by protein complex formation between the antibodies and AMPylated antigens on size exclusion chromatography. In order to determine the detection limits of recognition (Figure 2A), we tested all three antibodies in WB on a dilution series of recombinant Cdc42-Tyr-AMP, -Thr-AMP, and Rab1-Tyr-AMP, respectively. They all showed similar performance on all targets and modified side chains: All three antibodies were able to recognize up to as little as 2 ng or even lower amounts of AMPylated protein. Antibody 1G11 detected modified Cdc42 at the Thr side chain with less sensitivity than at the Tyr side chain, and 7C11 detected Tyr modified GTPases with less sensitivity than Thr modified protein. Antibody 17G6 did not show a preference for a specific AMPylated side chain.

**Figure 2:**
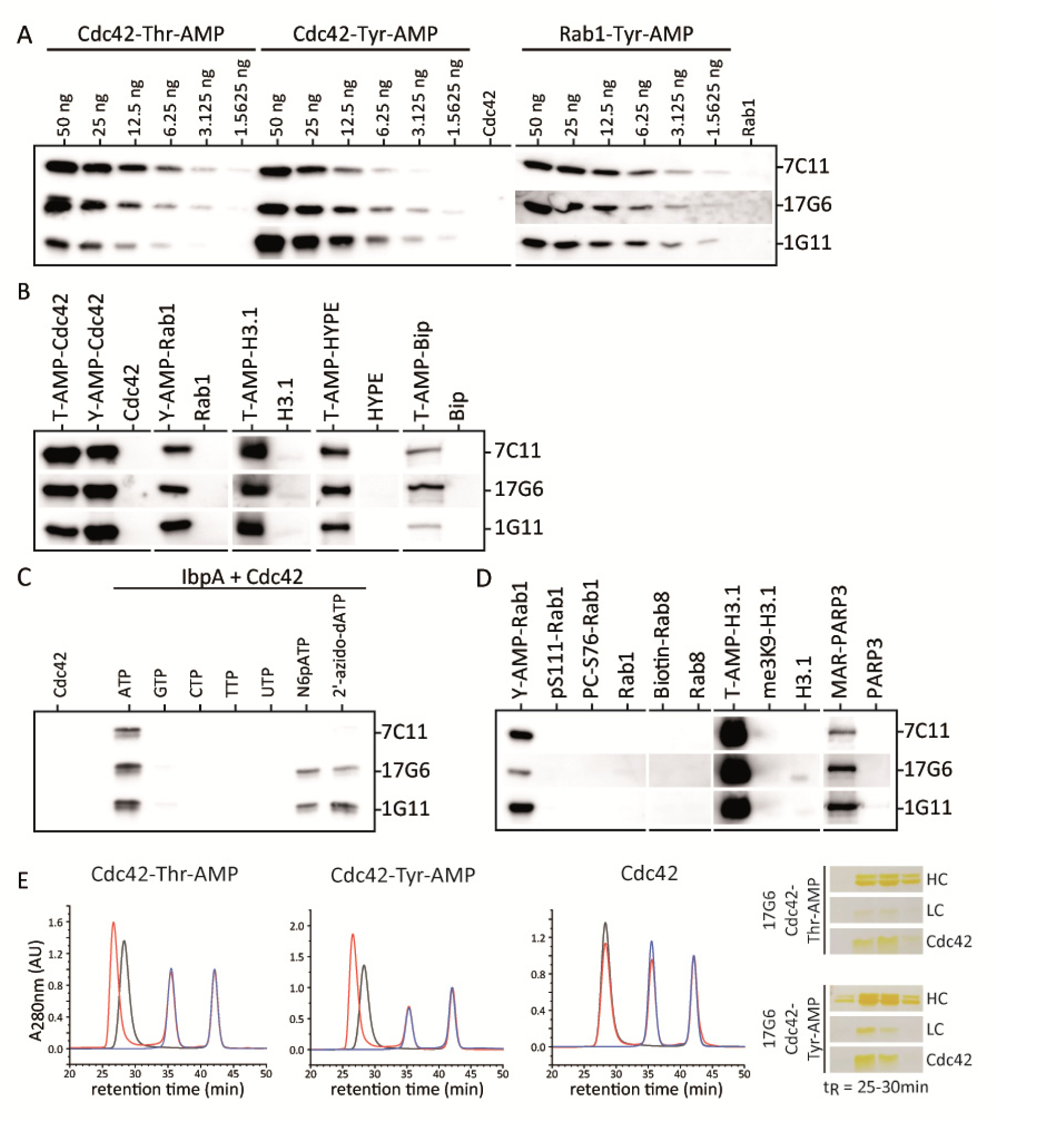
Generated anti-AMP-antibodies are highly specific for AMPylation. **A** Detection limits of AMPylated protein by the monoclonal anti-AMP antibodies in WB. Dilutions rows starting from 50 ng recombinant Cdc42-Thr-AMP, -Tyr-AMP and Rab1-Tyr-AMP, respectively, were analyzed in WB using all three monoclonal anti-AMP antibodies as indicated. **B** Broadness of AMPylated target recognition by the monoclonal anti-AMP antibodies in WB. 50 ng recombinant protein as indicated were analyzed in WB using all three monoclonal anti-AMP antibodies as indicated. **C** Evaluation of specificity towards AMPylation by the monoclonal anti-AMP antibodies in WB. IbpA was employed due to its ability to incorporate all indicated nucleotides into Cdc42 as NMPylation. 50 ng recombinant Cdc42 modified with nucleotides as indicated were analyzed in WB using all three monoclonal anti-AMP antibodies as indicated. **D** Cross-reactivity with other PTMs by the monoclonal anti-AMP antibodies in WB was analyzed by blotting 50 ng of recombinant protein as indicated. All three monoclonal anti-AMP antibodies cross reacted with mono-ADP-Ribosylation (MAR) on PARP3. **E** Native binding of AMPylated Cdc42 by monoclonal anti-AMP antibody 17G6 analyzed by analytical SEC. 40 µg antigen were mixed with 60 µg antibody, including 50 µM Vitamin B12 as internal standard. In black antibody 17G6 alone, in blue antigen alone as indicated, in red co-incubation of antibody 17G6 and antigen as indicated. Shifted antibody peaks (red) were fractionated and analyzed for co-elution of antibody and antigen by silver stained SDS PAGE.

Next, we aimed to confirm backbone independent recognition of AMPylation by applying all three antibodies on a broad range of AMPylated targets (Figure 2B), and indeed, the recognition of AMPylation was not limited to small GTPases. In addition, AMPylated Hype, Bip, and H3 were also recognized representing very different protein classes, sizes and folds. It therefore seems likely that other proteins will be recognized as well, which is a crucial prerequisite for target identification. Native binding of the antibodies to their modified antigens was investigated by complex formation with different AMPylated proteins using size exclusion chromatography (Figure 2E). The shift of retention time of the antibody peaks upon incubation with AMPylated proteins towards higher molecular weight but no shift for incubation with non-modified counterparts for all antibodies illustrates strong and specific binding of modified targets. The shifted antibody peaks were further collected by fractionation and analyzed by SDS PAGE for their co-elution with the antigens. Indeed, in case of AMPylated antigens, the antibodies co-eluted with their antigens. In this experiment, the same preferences for side chains were observable as already deduced from studies by WB (denaturing conditions): Antibody 1G11 shows a preference for AMPylated tyrosine, exemplified by a striking peak shift upon Rab1-Tyr-AMP binding but little shift for Cdc42-Thr-AMP. Antibody 17G6 shows broad recognition of all modified targets, whereas 7C11 prefers threonine AMPylation and does not show binding of Rab1-Tyr-AMP (Supp. Figure S2). Notably, Rab1-Tyr-AMP appears to be a difficult antigen for native as well as denatured recognition by the new antibodies: Already during selection, Rab1-Tyr-AMP recognition in WB was one of the main hurdles for most candidates, and there were only few candidates who showed a strong signal in WB.

To test the antibodies’ specificity towards the transferred nucleotide and their ability to differentiate AMPylation from e.g. GMPylation, recombinant IbpA was used to introduce UMPylation, GMPylation, CMPylation and TMPylation onto Cdc42 (Figure 2C). In addition, the recognition of two reactive ATP analogs that have been previously described in the context of AMPylation, N6-Propargyl-ATP (N6pATP) (Grammel et al., 2011; Yu et al., 2014) and 2’-Azido-2’-dATP (Wang & Silverman, 2016), was tested. All antibodies successfully differentiated between the nucleotides and specifically recognized AMPylation in Cdc42. Using ATP analogs instead of ATP, we could confirm that the antibodies are also sensitive to base and ribose modifications, and only antibodies 17G6 and 1G11 showed slight recognition of modified ATP-analogs (Figure 2C). This preference for AMPylation suggests a recognition of the adenine ring system by the antibodies.

After proving that the antibodies are sensitive and specific for AMPylation, we tested various other common PTMs for their ability to cross-react with the antibodies, to rule out false positive signals from competing modifications (Figure 2D). We tested phosphorylated (pS111) and phosphocholinated (PC-S76) Rab1b in direct comparison to its AMPylation, as well as biotinylated Rab8, trimethylated (me3K9) H3.1 in direct comparison to its AMPylation, and mono-ADP-ribosylated (MARylated) PARP3. To our satisfaction, the antibodies did not recognize phosphorylation, phosphocholination, biotinylation, or trimethylation on the chosen example proteins. However, the antibodies cross reacted with MARylation on PARP3, most likely recognizing the present adenosine moiety.

### Anti-AMP-antibodies can shift bias between AMPylation and MARylation

Our findings show that the developed anti-AMP-antibodies also detect mono-ADP-ribosylation as exemplified by auto-modified PARP3 (Figure 2D). We therefore screened different additives to the primary antibody incubation step during WB for their ability to abrogate reactivity with ADP-ribosylation, while keeping recognition of AMPylation intact (Figure 3A). Adenine, AMP, ADP, ATP, ADP-Ribose (ADPR) and nicotinamide adenine dinucleotide (NAD^+^) were selected for their similarity to both modifications and their potential ability to compete with binding of ADP-ribosylation or AMPylation and displace modified proteins. MnCl_2_ and MgCl_2_ as divalent cations were chosen for their ability to possibly complex the negatively charged diphosphate present in ADP-ribosylation but not AMPylation, thereby shielding negative charge that could potentially be relevant for antibody binding. Hydroxylamine treatment of the membrane after blotting reportedly results in specific cleavage of ADP-ribosylation at aspartate and glutamate side chains (Moss et al., 1983), but has not be previously tested regarding the stability of AMPylation. None of the tested additives were able to selectively reduce reactivity towards neither AMPylation nor MARylation, without significantly reducing overall sensitivity of the antibodies at the same time. Nevertheless, AMP, ADP, ATP and NAD^+^ were able to reduce AMPylation signals to some extent, while the MARylation signal remained largely unaffected. However, keeping in mind that PARP3 has 14 reported auto-MARylation sites (Vyas et al., 2014), whereas Cdc42 is single AMPylated, this loss of signal in AMPylation but not MARylation might be due to the multiple modifications on MAR-PARP3 and therefore not be easily translatable towards other single ADP-ribosylated proteins, where these additives might also result in signal loss. As expected, hydroxylamine treatment resulted in a strong loss of MARylation signal due to cleavage of the ADP-ribosyl group. By contrast, the AMPylation signal remained entirely unaffected. The residual signal of auto-modified PARP3 is most likely resulting from the two reported auto modification sites at lysin6 and lysin37 (Vyas et al., 2014) that will not be cleaved by hydroxylamine.

**Figure 3:**
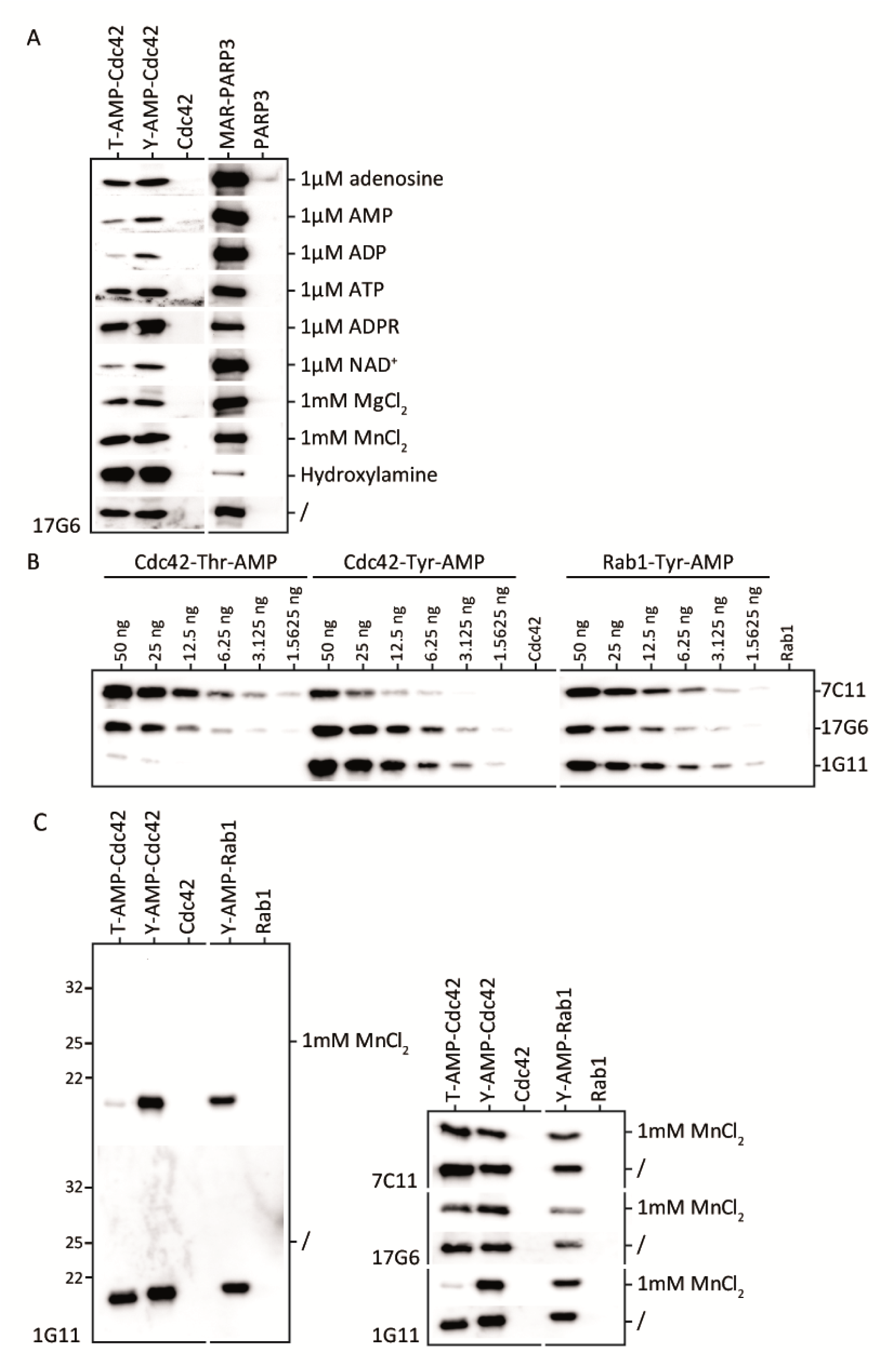
Anti-AMP-antibodies can shift bias between detection of AMPylation and MARylation. **A** Recognition of AMPylation vs. MARylation by antibody 17G6 can be fine-tuned using additives as indicated during primary antibody incubation or hydroxylamine treatment. 50 ng recombinant Cdc42-Thr-AMP, -Tyr-AMP as well as mono-ADP-ribosylated PARP3 as indicated were analyzed in WB. Additives as indicated, with the exception of hydroxylamine, were added during incubation with primary anti-AMP antibody. Hydroxylamine treatment to cleave off ADP-ribosylation at Asp and Glu took place for blotting before primary antibody incubation. **B** Detection limits under the influence of 1 mM MnCl_2_. Dilutions rows starting from 50 ng recombinant Cdc42-Thr-AMP, -Tyr-AMP and Rab1-Tyr-AMP, respectively, were analyzed in WB using all three monoclonal anti-AMP antibodies as indicated in the presence of 1 mM MnCl_2_. **C** Addition of 1 mM MnCl_2_ reduces antibody background and causes antibody 1G11 to regain tyrosine-AMP specificity. 50 ng recombinant protein as indicated were analyzed in WB using all three monoclonal anti-AMP antibodies as indicated.

The addition of 1 mM MnCl_2_ during the primary antibody incubation step, while not affecting signal intensity, resulted in a significantly reduced background. Strikingly, the addition of MnCl_2_ also resulted in a sharply enhanced tyrosine specificity for antibody 1G11 in the presence of MnCl_2_ (Figure 3C). The addition of MnCl_2_ was therefore further evaluated in regard to the previously tested detection limits of the antibodies. We confirmed that the detection limits of antibodies 17G6 and 7C11 towards AMPylated antigens was not significantly changed by addition of MnCl_2_, while 1G11’s ability to detect Thr-AMP-Cdc42 was greatly diminished (Figure 3B).

In summary, our newly developed antibodies are a combined tool for detection of AMPylation and mono-ADP-ribosylation. By addition of MnCl_2_ to the primary antibody incubation steps in WB, the background of the antibodies can be significantly reduced and 1G11 shows pronounced tyrosine specificity. By hydroxylamine treatment of membranes after blotting, glutamate and aspartate linked ADP-ribosylation can be cleaved while AMPylation remains unaffected. Therefore, the combination of all three antibodies with addition of MnCl_2_ and hydroxylamine treatment results in a tool-kit, which is able to sensitively detect ADP-ribosylation and AMPylation, to differentiate between the two, and in case of AMPylation to recognize not only targets in general but also to give information on their modified side chains.

### Generated anti-AMP-antibodies recognize diverse cellular AMPylation

After thorough characterization and evaluation of our produced antibodies on purified and mass spectrometry (MS) confirmed antigens, we next evaluated the antibody performance on cell lysates in known contexts under denatured as well as native conditions. The reproduction of previous results with these new tools is crucial for the trust in future findings and a smooth transition from previously used tools.

Previously, it could be shown that Bip AMPylation by HYPE is lost in cells upon stimulation of endoplasmic reticulum (ER) stress e.g. by thapsigargin (Ham et al., 2014; Preissler et al., 2015). Cycloheximide, in contrast, will stall protein production, therefore relieving the ER of protein load, causing a significant increase in Bip AMPylation. We reproduced these findings – representative for all our antibodies - with antibody 17G6 and MnCl_2_ as additive in Cho-K1 cells (Figure 4A + B) and could confirm these previously published results. In order to verify that the antibodies’ previously confirmed ability to bind native AMPylated protein would translate into a successful immunoprecipitation (IP), we applied the antibodies in the same context of Bip AMPylation (Figure 4B and 5C). Using our antibody 17G6, we successfully performed an immunoprecipitation of AMPylated Bip, first as recombinant protein (Figure 4B), and afterwards from thapsigargin and cycloheximide treated CHO-K1 cells (Figure 4C), respectively. Using recombinant Bip and Bip-AMP we can show that immunoprecipitation is dependent on the presence of antibody 17G6 and specific for AMPylation, the non-modified Bip is not precipitated. The detection of a successful IP from cell lysates was hampered by the detection limit of the anti-Bip antibody, which did not recognize Bip at less than 50 ng per lane. With detection limits of the anti-AMP antibodies far lower, the detected band by the anti-AMP antibody 17G6 in untreated CHO-K1 whole cell lysates (in WB, Figure 4A) might correspond to less than 5 ng in 20 µg loaded lysate, while the less sensitive anti-Bip antibody leads to a more pronounced signal. Therefore, we had to assume that the percentage of AMPylated Bip in the untreated CHO-K1 cell lysates was very low. Consequently, AMPylation was stimulated by cycloheximide treatment in order to create enough pulldown material for detection with anti-Bip antibody, resulting in a band intensity of Bip-AMP comparable to 50 ng in 20 µg loaded whole cell lysate using the anti-AMP antibody 17G6 (Figure 4C).

**Figure 4:**
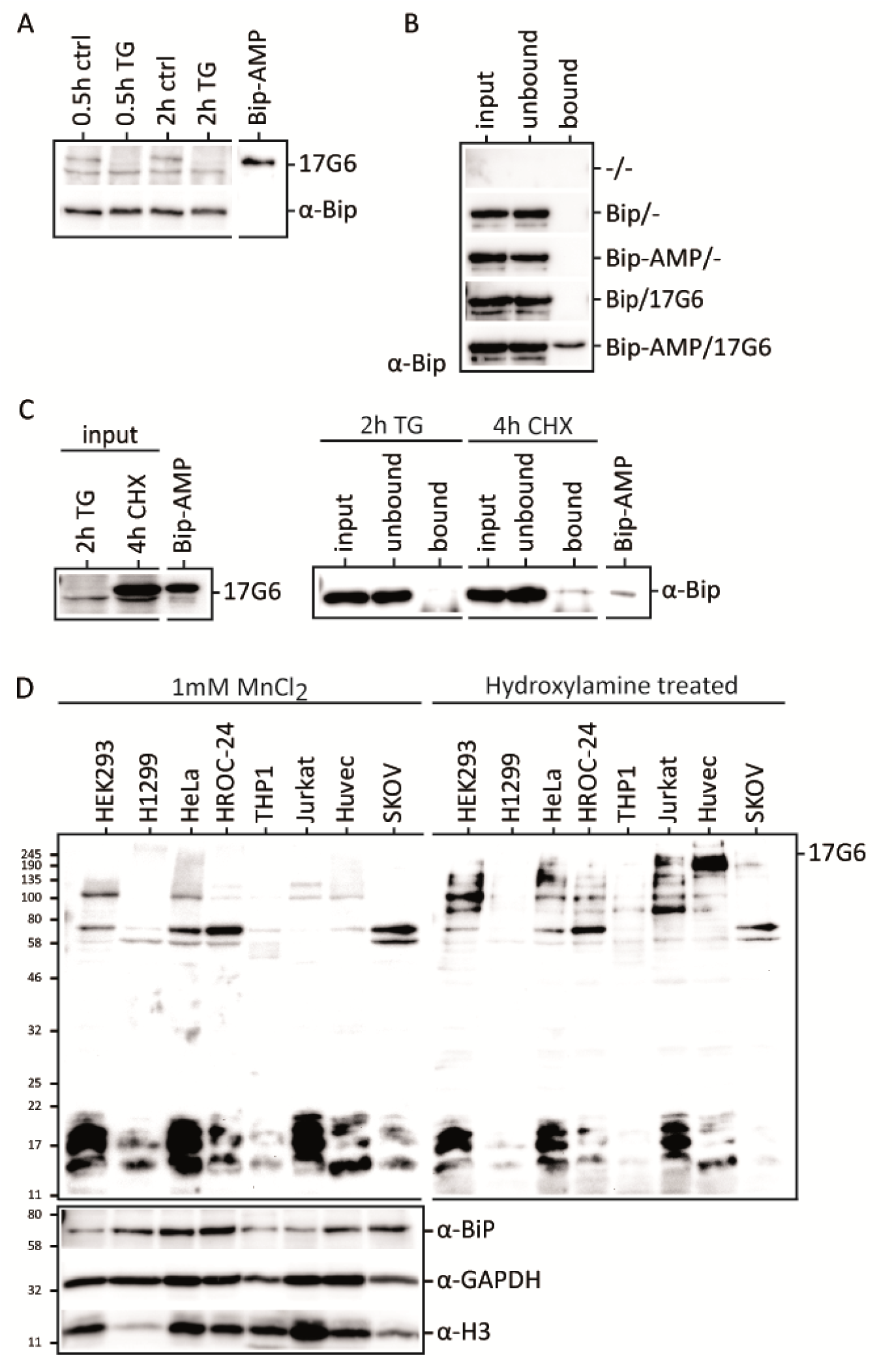
Generated anti-AMP-antibodies recognize diverse cellular AMPylation. **A** Reproduction of previously published data confirms loss of Bip-AMPylation upon ER stress by thapsigargin in WB. 20 µg treated (as indicated) ChoK1 cell lysate per lane or 50 ng recombinant Bip-AMP were analyzed in WB by antibody 17G6 and anti-Bip antibody. **B** Successful IP with antibody 17G6 on recombinant BiP-AMP confirms that antibody 17G6 is AMP-specific. **C** Successful IP of endogenous Bip-AMP with antibody 17G6 from treated (as indicated) ChoK1 cell lysates confirms applicability in immunoprecipitation. 50 ng recombinant Bip-AMP were blotted as control. **D** Using antibody 17G6 on various immortalized and cancer cell lines reveals diverse cellular AMPylation. 20 µg cell lysate per lane as indicated were blotted and probed with antibody 17G6 using 1 mM MnCl_2_ as additive. Afterwards cells were treated with 1 M hydroxylamine to cleave ADP-ribosylation at aspartate and glutamate residues and reprobed with antibody 17G6 using 1 mM MnCl_2_. Antibodies against Bip, GAPDH and Histone H3 serve as loading control.

The successful immunoprecipitation of endogenous amounts of AMPylated Bip from cell lysates shows potential for future target identification by IP, especially in combination with MS.

After the thorough characterization of our antibodies’ performance in WB, we asked the question whether these sensitive tools were able to detect new AMPylation bands in cell lysates. We therefore screened a number of available immortalized and cancer cell lines for occurrence of AMPylation bands using our newly developed monoclonal antibodies (Figure 4D and Supplemental Figure S4). And indeed, our anti-AMP antibodies were able to detect a multitude of bands in the range of 58-245 kDa and 11-22 kDa. Strikingly, some bands differed among cell lysates, while others were distinctive and reoccurring. While some cell lines such as THP-1 show very little to no AMPylation signals, other cell lines such as HeLa, HEK293 and Jurkat cells show strong AMPylation signals, especially in the region of 11-22 kDa. Treatment of membranes with hydroxylamine to cleave off ADP-ribosylation at aspartate and glutamate residues does not significantly diminish these bands. Furthermore, these bands are not detected by the anti-pan-ADPR binding reagent (Merck), or tyrosine specific anti-AMP antibody 1G11 (see Figure S4), but both anti-AMP antibodies 7C11 and 17G6, strongly suggesting AMPylation at threonine residues. Another reoccurring band at 70 kDa, most likely representing Bip-AMP, is strongly differing in intensity among cell lysates: While H1299, THP-1, Jurkat and Huvec cells do not show this band at all, it is strongly represented in HeLa, HROC-24 and SKOV cells.

## Discussion

Here, we report and characterize three new monoclonal anti-AMP antibodies recognizing AMPylation independent of the protein backbone. In order to reduce the inherent batch to batch variability of the previously published polyclonal antibodies, as well as generate defined specificities, we created monoclonal antibodies in mice. The reproducibility crisis of antibodies in recent years (Baker, 2015), as well as the limitations of commercially available anti-AMPylation-antibodies (Hao et al., 2011) (Sigma-Aldrich ABS184 and 09-890) let us to perform a thorough evaluation of the new monoclonal antibodies’ performance in two different applications. For denatured recognition, we tested sensitivity, specificity, and cross reactivity of our antibodies in Western Blots. For native recognition, we studied complex formation of the antibody with different modified targets by size exclusion chromatography, as well as confirmed native binding in an immunoprecipitation experiment. The antibodies were generated with the help of an AMPylated synthetic peptide with reduced backbone complexity. A major bottle neck in antibody generation based on synthesized peptides, which is also reported for the generation of other anti-PTM antibodies such as anti-phospho antibodies (Archuleta et al., 2011), is the phenomenon of predominantly positive peptide ELISA readings against modified hapten, that do not translate to a positive WB performance. Common procedure is to only select via ELISA against the modified hapten. According to Archuleta et al. (Archuleta et al., 2011), this method selects antibodies, whose performance fails in other applications in 25-50% of cases. However, in our selection process we observed a high correlation between positive ELISA readings against modified protein, which we performed in addition to peptide ELISA, and good WB performance. The inclusion of a native AMPylated protein in form of Cdc42-Thr-AMP in the ELISA screening process allowed us to generate monoclonal antibodies combined with efficient preselection of candidates before WB evaluation. We therefore recommend including native modified protein in the ELISA screening process for all anti-PTM-antibodies.

First efforts in the creation of anti-AMP antibodies were undertaken in 1984 by fusing AMP directly to the carrier protein BSA (Chung & Rhee, 1984), thus generating murine monoclonal antibodies that were purified from ascitic fluid and employed in the purification of AMPylated glutamine synthetase. Later on, other antibodies were accidentally produced by aiming for ADP-ribose antibodies, where the hapten was degraded to contain AMP, resulting in antibodies recognizing free 5’-AMP (Bredehorst et al., 1978; Meyer & Hilz, 1986). Hao et al. (Hao et al., 2011) achieved polyclonal antibodies by immunization of rabbits with a synthetic seven amino acid long Rac1-peptide containing a threonine AMPylation (now commercially available as Anti-pan-AMPylated Threonine Antibody 09-890, Sigma-Aldrich Merck). After depletion with tyrosine-AMPylated protein the serum was reported to detect threonine AMPylation independently of protein backbone and structure, in WB as well as IP. The most recent antibody was produced by an AMPylated Rab1b peptide of 13 amino acids coupled to KLH in rabbit, resulting in polyclonal serum, aided again by efficient synthesis of AMPylated peptides (Smit et al., 2011). However, both published rabbit antibodies are hampered by low sensitivity, and little characterization, especially concerning cross reactivity with other PTMs and recognition of targets outside the protein class of small GTPases or Bip is published. In addition, all recently developed antibodies are polyclonal, with the accompanying challenges of batch to batch reproducibility and reliability of tool development on basis of that antibody. Considering the special challenges connected with the generation of antibodies that target PTMs (Hattori & Koide, 2018), and their necessity for extensive characterization, polyclonal antibodies are not an ideal choice. A stringent retesting of every new batch regarding proper AMP-recognition and lack of cross reactivity would have to be performed before application to cell lysates. Previous antibodies therefore represented no general recognition tool of AMPylation, especially if searching for new targets and effects, where the number of potential false negative or false positive findings would render them unreliable. Our experiments show that all commercially available anti-AMP antibodies offer no broad recognition of targets, despite claiming to recognize AMPylation backbone independently, and are exhibiting a significant amount of false positive and negative reactions in our in-house testing. The limitations in performance and cross reactivity of both anti-ADPR-reagents and anti-AMPylation antibodies in combination causes the danger of false positives for ADP-ribosylation as well as false negatives in AMPylated proteins, and a bias in AMPylation research towards small GTPases and threonine modifications (Figure 1A). As many researchers lack suitable positive and negative controls of protein of interest, these performance failures might never be detected.

Little is known about AMPylation in eukaryotic cells outside the modification of Bip in the context of ER stress. In accordance with recent publications (Kielkowski et al., 2020; Sreelatha et al., 2018), the application of our new monoclonal antibodies to cell lysates of immortalized and cancer cell lines hint at a much stronger prevalence of AMPylation than perceived for a long time. The limited number of tools, especially in medium to high throughput, has hampered reliable detection of AMPylation in cellular systems. Our antibodies expand the available toolbox by offering sensitive detection and enrichment of AMPylation, at the same time requiring little resources that might hamper applicability in a standard laboratory. They therefore open new opportunities in an expanding research field. The recent antibody “reproducibility crisis” especially in regards to antibodies targeting PTMs (Baker, 2015; Egelhofer et al., 2011) suggested a thorough characterization of the AMPylation-specificity and sensitivity of our new monoclonal antibodies in the applications WB, ELISA and IP. With their high sensitivity and broad target recognition, they overcome the limitations of previously published anti-AMP antibodies and create opportunities for new target identification and study of cellular AMPylation. Our data suggest that they can successfully be used for enrichment of AMPylated proteins and peptides for mass spectrometry to overcome the limitation of low occurrence of AMPylation in proteomic studies. As all three monoclonal antibodies are sequenced, thereby enabling recombinant antibody production, they form a good basis for long-term reproducibility in AMPylation research.

## Materials and Methods

### Solid phase peptide synthesis (SPPS)

The peptides S2 and S3 (Supplemental Figure S5) were synthesized on a MultiSyntech Syro I automated peptide synthesizer, using Tentagel Rink-amide resin as solid phase employing 8 equivalents of amino acid and 7.8 equivalents HBTU, 7.8 equivalents HOBt and 16 equivalents DIPEA in DMF. Threonine-AMP building block S1 (Smit et al., 2011) was manually coupled according to below. The resin was pre-swollen by treatment with DCM (15 min). Removal of Fmoc protecting group was performed by treatment with 2 times 3 minutes followed by 1 time 9 minutes 20 vol% piperidine in DMF. The resin was washed with DMF three times. The building block S1 (1.7 equivalents) was coupled using 1.6 equivalents HATU, 1.6 equivalents HOAt and 3.5 equivalents DIPEA in DMF. The coupling reaction was allowed to proceed for 4 h at room temperature. N-terminal acetyl-capping was achieved by adding 50 equivalents acetic anhydride and 50 equivalents DIPEA in NMP to the resin (1 h), followed by washing with DMF. Before cleavage, the resin was washed thoroughly with DCM (5 times), isopropanol (5 times) and diethyl ether (5 times). The peptides were cleaved with 5% TIPS and 5% water in TFA (1 × 2 h + 2 × 10 min). An additional 10% water was added to the combined TFA-fractions and allowed to age for one hour, in order to ensure complete hydrolysis of the acetonide moiety. The cleavage mixture was evaporated to dryness *in vacuo*. The resulting solid was dissolved in a minimum TFA and precipitated by the addition of 10 ml ice cold diethyl ether. The precipitate was dissolved in water/acetonitrile and subsequently lyophilized overnight. Pure peptides were obtained after purification by preparative HPLC. Preparative HPLC purifications of the peptides were performed with an Agilent 1260 Infinity series instrument equipped with a Phenomenex Luna (5 µm, C18(2) 100 Å, 250 × 21.2 mm) column. Used mobile phases were water with 0.1% TFA (eluent A) and acetonitrile with 0.1% TFA (eluent B). Analytical HRMS-HPLC was performed on an Agilent 1290 Infinity II series instrument equipped with an Agilent Extend (1.8 µm, C18, 100 Å, 50 × 2.1 mm) column and connected to an Agilent 6230 TOF LC/MS instrument. Used mobile phases were water with 0.1% FA (eluent A) and acetonitrile with 0.1% FA (eluent B). For complete description see supplement.

### Immunogen preparation

Immunogen peptide S2 (ACNH-CGAGT(AMP)GAG-NH2) was conjugated with KLH and BSA as immunogen via its N-terminal cysteine (GenScript).

### Immunization

3 BALB/c and 2 C57 BL/6 mice were immunized with either S2 conjugated to KLH (group A, BALB/c mice #8534 - #8536 and C57 BL/6 mice #8537 - #8538) or BSA (group B, BALB/c mice #8539 - #8541 and C57 BL/6 mice #8542 - #8543), respectively, according to the conventional protocol of GenScript (Table 2), resulting in 10 immunized mice in total.

**Table 1:** Strategy for monoclonal anti-AMP antibody generation in mice. Experiments were conducted as indicated as a service by GenScript, Piscataway Township, New Jersey.

| Stage | Description | Shipment |
| --- | --- | --- |
| Immunogen preparation | Peptide conjugation with KLH and BSA. | - |
| Phase I: Immunization | Group A: 3 BALB/c +2 C57 mice, immunization with KLH conjugates, Group B: 3 BALB/c +2 C57 mice, immunization with BSA conjugates, Immunization with conventional protocol, indirect ELISA primary screening with target peptide (AcNH-CGAGT(AMP)GAG-NH<sub>2</sub>), and confirmatory screening by indirect ELISA by target protein (Cdc42-AMP) | 15µl sera of 10 animals |
| Phase II: Cell fusion | 2 cell fusions, indirect ELISA primary screening with target peptide (AcNH-CGAGT(AMP)GAG-NH<sub>2</sub>), and confirmatory screening by indirect ELISA by target protein (Cdc42-AMP) | Supernatants of 10 clones,<br>2ml per clone |
| Phase III: Subcloning, Screening and Expansion | 3 cell lines subcloning, indirect ELISA primary screening with target peptide (AcNH-CGAGT(AMP)GAG-NH<sub>2</sub>), and confirmatory screening by indirect ELISA by target protein (Cdc42-AMP) | 2 vials and 5ml supernatant per subclone |
| Antibody production | 1 L roller bottle cell culture and protein A purification, indirect ELISA primary screening with target peptide (AcNH-CGAGT(AMP)GAG-NH<sub>2</sub>), and confirmatory screening by indirect ELISA by target protein (Cdc42-AMP) | >15mg purified antibody per subclone |

**Table 2:** Immunization schedule.

| Procedure | Schedule | Dosage and route | Adjuvant |
| --- | --- | --- | --- |
| Pre-Immune Bleed | T= - 4 days |  |  |
| Primary Immunization | T= 0 days | 50 μg /animal, i.p. | CFA |
| Boost 1 | T= 14 days | 25 μg /animal, i.p. | IFA |
| Test Bleed 1 | T= 21 days |  |  |
| Boost 2 | T= 28 days | 25 μg /animal, i.p. | IFA |
| Test Bleed 2 | T= 35 days |  |  |
| Final Boost | T= 50±7 days | 25 μg /animal, i.p. | IFA |
| Cell Fusion | 4 days after final boost |  |  |

Test bleed: 7 days after each boost immunization, immunized animal sera were tested by indirect ELISA and competitive ELISA for immune response by GenScript. Western Blot evaluation of pre-sera and sera after 3rd immunization against 200 ng of purified protein/lane using a 1:1000 dilution was performed in-house as described. Animals #8538 (group A) and #8542 (group B), both C57 BL/6 mice, were selected for phase II.

### Phase II: cell fusion and screening

For cell fusion and clone plating, two fusions were performed by electro-fusion. The average fusion efficiency at GenScript is around 1 hybridoma/2,000 splenocytes, thus the anticipated hybridoma clones would be ∼ 2×10^4^. All fused cells from each cell fusion were plated into 96-well plates. Up to 15 plates were used for each fusion. For the primary binder screening, the conditioned medium was screened by ELISA with the target peptide. In the confirmatory screening, the conditioned medium of primary binder screening positive clones were screened against the positive screening material (Cdc42-Thr-AMP) and counter screening material (Cdc42). The expected clones should be positive against target peptides, positive screening material while negative against the negative peptide and counter screening material. For clone expansion and freezing, 10 positive clones were expanded into 24-well plates. 2 ml of supernatant (conditioned media) were collected for each clone and cells were frozen down to avoid clone loss (GenScript). The conditioned media of all 10 positive clones were analyzed in-house by WB against 200 ng of purified protein/lane using a 1:10 dilution as described. Clones 17G6 (#8542), 1G11 and 7C11 (#8538) were selected for phase III.

### Phase III: subcloning, screening, expansion

For subcloning, 3 positive primary clones were sub-cloned by limiting dilution to ensure that the sub-clones were derived from a single parental cell. The clones were carried for a maximum of 3 generations. Subcloning was screened by ELISA as before. For monoclone cryopreservation, two stable sub-clonal cell lines of each parental clone were chosen for cryopreservation based on the result of ELISA (GenScript). Positive cell supernatants were evaluated by WB against 200 ng of purified protein/lane using a 1:10 dilution in-house as described. The stable sub-clonal cell lines 17G6-1 (isotype IgG2b, k), 1G11-F3-3 (isotype IgG2b, k) and 7C11-1 (isotype IgG2a, k) were chosen for production and isotyped, and the cell lines stored with 2 vials at GenScript and 2 vials in-house. They were negatively tested for mycoplasma, detected by the PCR Mycoplasma Detection Set (TAKARA BIO INC, Kusatsu, Japan).

### Antibody production

Three stable sub-clonal cell lines were each cultured in 1 l roller bottle cell culture using SFM + 2 % low IgG FBS culture medium. Monoclonal antibody was Protein A purified from the supernatant and stored in phosphate buffered saline (PBS, pH 7.4) with 0.02 % sodium azide as preservative. Purity was measured by SDS-PAGE and concentration by NanoDrop Spectrophotometer A280nm (Thermo Fisher Scientific Inc., Waltham, Massachusetts) (GenScript). This way 36.38 mg of 7C11 with 95% purity, 17.10 mg 17G6 with 91 % purity and 30.99 mg 1G11 with 92% purity were produced.

### Molecular biology

Unless otherwise indicated, all genes were codon optimized for expression in *E. coli* by omitting rare amino acid codons, and all cloning was done by sequence and ligation independent cloning (SLIC) using T4 DNA polymerase (New England Biolabs, Ipswich, Massachusetts).

The human Cdc42 1-179aa Q61L (referred to as Cdc42)-encoding DNA was cloned into a modified pGEX-4T-1 vector (GE Healthcare, Chicago, Illinois) as previously described (Barthelmes et al., 2020), resulting in a construct with a N-terminal glutathione S-transferase (GST) tag followed by the Tobacco Etch Virus (TEV) protease cleavage site. As previously described (Schoebel et al., 2009), the human Rab1b 3-174aa (referred to as Rab1b)-encoding DNA was cloned into a modified pMAL vector (New England Biolabs), resulting in a construct with a N-terminal hexahistidine (6xHis) tag, followed by maltose-binding protein (MBP) and the TEV protease cleavage site. The human Bip 19-654aa (referred to as Bip)-encoding DNA was cloned into a modified pProEx™-HTb vector (Thermo Fisher Scientific Inc.), resulting in a construct with a N-terminal 6xHis tag, followed by the TEV protease cleavage site. The human Hype 102-458aa E234G (referred to as Hype)-encoding DNA was cloned into a modified pMAL vector (New England Biolabs), resulting in a construct with a N-terminal 6xHis tag, followed by the HaloTag^®^, the TEV protease cleavage site and a Strep-tag^®^ II. The *Vibrio parahaemolyticus* VopS 31-387aa (referred to as VopS)-encoding DNA was cloned into a modified pMAL vector (New England Biolabs) as previously described (Barthelmes et al., 2020), resulting in a construct with a N-terminal 6xHis tag, followed by MBP and the TEV protease cleavage site. The *Histophilus somni* IbpA 3483-3797aa I3455C (referred to as IbpA)-encoding DNA was cloned into a modified pSF vector (Oxford Genetics Ltd, Oxford, UK), resulting in a construct with a N-terminal decahistidine (10xHis) tag, followed by MBP, the TEV protease cleavage site and a 3xFLAG^®^ tag. The *Legionella pneumophila* DrrA 8-533aa (referred to as DrrA)-encoding DNA was cloned into a modified pET19 vector (Merck Millipore, Burlington, Massachusetts) as previously described (Müller et al., 2010), resulting in a construct with a N-terminal 6xHis tag and the TEV protease cleavage site. The *Legionella pneumophila* AnkX 1-800aa (referred to as AnkX)-encoding DNA, which previously had been amplified from *Legionella pneumophila* genomic DNA (Goody et al., 2012), was cloned into a modified pSF vector (Oxford Genetics) as previously described (Ernst et al., 2020), resulting in a construct with a N-terminal 10xHis tag, followed by enhanced green fluorescent protein (eGFP) and the TEV protease cleavage site. Human Rab8a 6-176aa (referred to as Rab8a)-encoding DNA was cloned into a pet51b(+) vector (Merck Millipore), resulting in a construct with a N-terminal Strep^®^ II tag and enterokinase cleavage site and a C-terminal 10xHis tag. All site-specific mutagenesis was performed with the Q5 Site-Directed Mutagenesis Kit (New England Biolabs).

### Recombinant expression and purification of proteins

Recombinant human histone H3.1 was purchased from New England Biolabs (M2503S), and active human PARP3 from Sigma-Aldrich, St. Louis, Missouri (SRP0194-10UG, Lot #8050330111). Human pSer111 Rab1b was a kind gift of Dr. Sophie Vieweg and was produced as published before (Vieweg et al., 2020).

Cdc42, VopS, Rab1b, DrrA and AnkX were expressed and purified as previously described (Barthelmes et al., 2020; Ernst et al., 2020; Müller et al., 2010; Schoebel et al., 2009). In brief, plasmids were transformed into chemically competent BL21 (DE3) cells (Cdc42, Rab1b, IbpA) or Lemo21 cells (VopS) or BL21-CodonPlus (DE3)-RIL cells (DrrA, Rab8a, AnkX) or Rosetta 2 cells (Bip, HYPE) and protein was expressed in LB medium after induction with 0.5 mM isopropyl-β-dithiogalactopyranoside (IPTG) for 20 h at 25 °C (Cdc42, Rab1b) or 20 °C (VopS, DrrA, Rab8a, AnkX, IbpA) or 23 °C (HYPE) or 3 h at 37 °C (Bip). Cells were harvested, washed with PBS and lysed in 50 mM 4-(2-hydroxyethyl)-1-piperazineethanesulfonic acid (Hepes) pH 7.5, 500 mM sodium chloride (NaCl), 1 mM MgCl_2_, 2 mM ß-mercaptoethanol (ßMe), 10 µM guanosine diphosphate (GDP), 1 mM PMSF (Cdc42, Rab1b, Rab8a) or 50 mM Tris pH 8.0, 500 mM NaCl, 5% (v/v) glycerol, 2 mM β-Me, 1 mM PMSF (AnkX) or 50mM Hepes pH 7.5, 500 mM NaCl, 1 mM MgCl_2_, 2mM ßMe, 1 mM PMSF (VopS, HYPE) or 50 mM Hepes pH 8.0, 500 mM lithium chloride (LiCl), 2 mM β-Me, 1 mM PMSF (DrrA, IbpA) or 50 mM Hepes pH 7.4, 400 mM NaCl, 20 mM imidazole, 1 mM PMSF (Bip) after addition of DNase I by French press at 1.8 kbar. The lysates were cleared by centrifugation.

For GST-tagged proteins (Cdc42), the lysate was loaded onto a pre-equilibrated GST-Trap column (GE) and eluted with 3-5 column volumes (CV) of the same buffer supplemented with 10 mM glutathione.

For His-tagged proteins (VopS, Rab1b, DrrA, Bip, HYPE, Rab8a, IbpA), the lysate was loaded onto a pre-equilibrated Ni^2+^-charged Bio-Scale Mini Nuvia IMAC Cartridge (Bio-Rad Laboratories, Hercules, California), washed with 30 mM imidazole and eluted with a fractioned gradient from 30 mM – 350 mM imidazole over 20 CV.

The protein containing eluate was digested with 6x-His tagged TEV during dialysis against 50 mM Hepes pH 7.5, 100 mM NaCl, 2 mM ßMe, 10 µM GDP (Cdc42, Rab1b) or 20 mM Hepes pH 8.0, 100 mM NaCl, 2 mM β-Me (DrrA, IpbA) or 20 mM Tris pH 8.0, 300 mM NaCl, 5% (v/v) glycerol, 2 mM β-Me (AnkX) 20 mM Hepes pH 7.4, 100 mM NaCl (Bip) or 20 mM Hepes pH 7.4, 100 mM NaCl, 1 mM MgCl_2_, 1 mM ßMe (HYPE) with a cut off of MW 6000-8000 (Serva) for 16h at 4°C.

Digested, formerly His-tagged proteins (Rab1b, DrrA, Bip, HYPE, AnkX, IbpA) were reapplied to the Ni^2+^-charged column pre-equilibrated with dialysis buffer in order to remove the His-tag, uncleaved protein and TEV protease. The flow through was collected, concentrated and applied to a HiLoad™ 16/600 Superdex™ 75pg column (GE Healthcare) in 20 mM Hepes pH 7.5, 50 mM NaCl, 1 mM MgCl_2_, 1 mM dithioerythritol (DTE), 10 µM GDP (Rab1b) or 20 mM Hepes pH 8.0, 100 mM NaCl, 2 mM DTE (DrrA) or 20 mM Tris pH 8.0, 300 mM NaCl, 5% (v/v) glycerol, 1 mM β-Me (AnkX) or 20 mM Hepes pH 7.4, 150 mM potassium chloride (KCl), 1 mM MgCl_2_, 1 mM tris(2-carboxyethyl)phosphine (TCEP), 5% glycerol (HYPE) or 20 mM Hepes pH 8.0, 100 mM NaCl, 1 mM MgCl_2_, 2 mM DTE (IbpA) or applied to a HiLoad™ 16/600 Superdex™ 200pg column (GE Healthcare) in 20 mM Hepes pH 7.4, 150 mM KCl, 10 mM MgCl_2_ (Bip). For VopS and Rab8a, protein containing eluate was concentrated and applied to a HiLoad™ 16/600 Superdex™ 75pg column (GE Healthcare) in 20 mM Hepes pH 7.5, 100 mM NaCl, 1 mM MgCl_2_, 1 mM DTT (VopS) or 20 mM Hepes pH 7.5, 50 mM NaCl, 1 mM MgCl_2_, 1 mM DTE, 10 µM GDP (Rab8a) without TEV digestion. For digested, formerly GST-tagged proteins (Cdc42), protein digestion was concentrated and applied to a HiLoad™ 16/600 Superdex™ 75pg column (GE Healthcare) in 20 mM Hepes pH 7.5, 50 mM NaCl, 1 mM MgCl_2_, 1 mM DTE, 10 µM GDP (Cdc42). During all steps of protein purification, fractions were collected and analyzed by Coomassie blue stained sodium dodecyl sulfate (SDS) polyacrylamide gel electrophoresis (PAGE). Fractions containing pure protein of interest were pooled, concentrated to around 10 mg/ml, snap-frozen in liquid nitrogen and stored in multiple aliquots at -80 °C.

### In vitro AMPylation of recombinant proteins

For Cdc42-Thr-AMP, 200 µM Cdc42 were incubated with 10 µM VopS in the presence of 800 µM ATP in 20 mM Hepes pH 7.5, 100 mM NaCl, 1 mM MgCl_2_, 1 mM DTE at 20 °C overnight. For Cdc42-Tyr-AMP, 10 µM Cdc42 were incubated with 0.1 µM IbpA in the presence of 1 mM ATP in 20 mM HEPES pH 7.4, 100 mM NaCl, 1 mM MgCl2, 1 mM TCEP, 10 µM GDP at 20 °C overnight. For Rab1-Tyr-AMP, 10 µM Rab1b were incubated in the presence of 50 µM ATP and 0.1 µM DrrA at 25°C as previously described (Müller et al., 2010). AMPylated Cdc42 and Rab1b were purified by size exclusion chromatography on a HiLoad™ 16/600 Superdex™ 75pg column (GE Healthcare) in 20 mM Hepes pH 7.5, 50 mM NaCl, 1 mM MgCl_2_, 1 mM DTE, 10 µM GDP and full AMPylation was confirmed by MS.

For H3-Thr-AMP and auto-AMPylated HYPE, 30 µM H3.1 were incubated with 25 µM HYPE in the presence of 10 mM ATP in 20 mM HEPES pH 7.5, 150 mM NaCl, 1 mM MgCl2, 2 mM DTT at 20 °C overnight. For Bip-Thr-AMP, 50 µM BiP were incubated with 2.5 µM HYPE in the presence of 1.5 mM ATP in 25 mM Hepes pH 7.4, 100 mM KCl, 4 mM MgCl_2_, 1 mM calcium chloride (CaCl_2_) for 2 h at 30 °C. Bip-AMP was purified with Protino™ Ni-NTA Agarose (Macherey-Nagel, Düren, Germany) in previously listed buffers according to the manufacturer’s instructions.

### In vitro NMPylation of recombinant Cdc42 by IbpA

10 µM Cdc42 were incubated with 0.1 µM IbpA in the presence of 0.5 mM of either CTP, UTP, TTP, GTP, N6pATP (Jena Bioscience, Jena, Germany) and 2’-Azido-2’-dATP (TriLink BioTechnologies, San Diego, California) in 20 mM Tris-HCl pH 7.4, 100 mM NaCl, 1 mM MgCl_2_, 1 mM TCEP, 10 µM GDP at 20 °C overnight. Successful NMPylation was confirmed by MS.

### In vitro auto-mono-ADP-ribosylation of PARP3

500 ng PARP-3 (Sigma-Aldrich) were incubated at 25°C in a 100 μL reaction volume in 20 mM HEPES pH 8.0, 5 mM MgCl2, 5 mM CaCl_2_, 0.01 % NP-40, 25 mM KCl, 1 mM DTT, 0.1 mg/mL salmon sperm DNA (Thermo Fisher Scientific), 0.1 mg/mL BSA (New England Biolabs) in the presence of 250 μM NAD+ for 30 minutes as published before (Gibson et al., 2017). The reaction was stopped by the addition of 5x SDS-PAGE Loading Buffer, followed by heating to 95°C for 5 min.

### In vitro phosphocholination or Rab1b by AnkX

10 µM Rab1b was incubated with 0.1 µM AnkX in the presence of 1 mM CDP-choline (Enzo Life Sciences, Farmingdale, New York) for 2 h at 23 °C as published before (Goody et al., 2012).

### In vitro biotinylation of Rab8a

200 mM EZ-Link^®^ Maleimide-PEG2-Biotin (Thermo Fisher Scientific) stock solution in DMSO was diluted 1:10 in 1x PBS to a final concentration of 20 mM Maleimide-PEG2-Biotin label. 200 µM Maleimide-PEG2-Biotin label were added to 100 μM of Rab8a in PBS for 2 h on ice, before Rab8a was washed 3 times with PBS in an Amicon filter (Merck Millipore, 10 kDa NMWL). Incorporation of label was confirmed by MS.

### Analytical size exclusion chromatography (aSEC)

In 100 µl, 40 µg antigen were mixed with 60 µg antibody, including 50 µM Vitamin B12 as internal standard. 90 µl sample were injected onto a Superdeep 10/300 200pg column (GE Healthcare) coupled to a Prominence HPLC system (Shimadzu, Kyōto, Japan) and run at 0.5 ml/min for 60 min in 20 mM HEPES pH 7.5, 150 mM NaCl. Protein retention times were detected at 280 nm (A280nm), and intensities were normed to Vitamin B12. Peaks containing antigen:antibody complexes were collected in 500 µl fractions. Fractions were supplemented with 6x Laemmli and concentrated in a SpeedVac alpha RVC (Martin Christ Gefriertrocknungsanlagen GmbH, Osterode am Harz, Germany) without heat to 200 µl. 10 µl concentrated fractions were analyzed by 15% SDS PAGE and silver stained.

### Mass spectrometry

To verify the degree of modification, samples containing 100 ng recombinant protein were run over a 5 μm Jupiter C4 300Å LC column (Phenomenex, Torrance, California) using the 1260 Infinity LC system (Agilent Technologies, Santa Clara, California) and then subjected to mass spectrometry with the 6100 Quadrupole LC/MS System (Agilent Technologies). The resulting ion spectra were deconvoluted using the Magic Transformer (MagTran) software (Zhang & Marshall, 1998).

### Western Blotting

200 ng or 50 ng recombinant protein as indicated or 20 µg cell lysate, respectively, were separated by SDS-PAGE and protein was transferred to MeOH-activated Immobilon^®^-P membrane (Merck Millipore) using Whatman paper and a transfer buffer of 48 mM Tris, 39 mM glycine, 1.3 mM SDS, 20% methanol. For the blotting procedure, a constant current of 0.7 mA/cm^2^ was applied to the V20-SDP semi-dry blotter (SCIE-PLAS, Cambourne, UK) for 2 h. After blotting, the PVDF membrane was blocked with Roti^®^-Block (Carl Roth, Karlsruhe, Germany) in Tris-buffered saline containing 0.1% Tween20 (TBS-T) for 1 h. Subsequently, the primary antibody was added to the blocking solution and incubated overnight at 4°C. Afterwards, the membrane was washed three times with TBS-T for 10 minutes and then incubated with a secondary antibody-peroxidase conjugate in TBS-T for 1 h. Again, the membrane was washed in TBS-T three times for 10 min, before the peroxidase signal was developed with the SuperSignal™ West Dura (Thermo Fisher Scientific) and chemoluminescence was detected using the Intas ECL Chemocam (Intas Science Imaging Instruments, Göttingen, Germany). Antibodies: Mouse pre-immune serum and antiserum after 3^rd^ immunization (GenScript) was used 1:1000. Cell supernatant from hybridoma clones and subclones (GenScript) was used 1:10. Purified monoclonal mouse anti-AMP antibodies 17G6, 1G11, 7C11 (GenScript) were used 1:1000 at 0.5 µg/ml. Monoclonal mouse anti-GAPDH sc-47724 (Santa Cruz Biotechnology, Dallas, Texas) was used 1:1000. Polyclonal rabbit anti-histone H3 antibody ab1791 (Abcam, Cambridge, UK) was used 1:5000. Polyclonal rabbit anti-AMPylated Tyrosine Antibody ABS184 (Merck Millipore) was used 1:1000. Polyclonal rabbit anti-pan-AMPylated Threonine Antibody 09-890 (Sigma-Aldrich) was used 1:2000. Recombinant rabbit anti-pan-ADP-ribose binding reagent MABE1016 (Merck Millipore) was used 1:1000. Polyclonal rabbit anti-GRP78/Bip antibody PA5-34941 (Thermo Fisher Scientific) was used 1:5000. Secondary goat anti-mouse lgG (H + L) HRP conjugate (Thermo Fisher Scientific) was used 1:20000. Secondary goat anti-rabbit IgG HRP (Sigma-Aldrich) was used 1:40000. Additives as indicated were added during the primary antibody incubation step at a final concentration of 1 µM for adenosine (Jena Bioscience), AMP (Sigma-Aldrich), ADP (Biosynth Carbosynth, Staad, Switzerland), ATP (Biosynth Carbosynth), ADPR (Sigma-Aldrich), NAD^+^ (Biosynth Carbosynth) or 1 mM for MnCl_2_, MgCl_2_ (VWR International, Radnor, Pennsylvania) respectively. Hydroxylamine treatment was performed as previously described (Gibson et al., 2017). In short, after development of membrane with anti-AMP antibodies, the membrane was incubated with 1 M hydroxylamine (Sigma-Aldrich) in blocking solution for 8 h at room temperature, washed three times in TBS-T, blocked again for 1 h at room temperature, and proceeded with a second round of anti-AMP antibody probing. All Western Blots on recombinant proteins were performed as technical duplicates. Analysis of cell lysate samples was performed as biological duplicate. Representative blots are shown.

### Cell culture

Chok-K1 FlpIn cells (Thermo Fisher Scientific) were cultured in RPMI-1640 medium (Sigma-Aldrich) supplemented with 10% FBS (Thermo Fisher Scientific). 90% confluent cells were stimulated by either 0.5 µM thapsigargin (Biosynth Carbosynth) for 2 h or 100 µg/ml cycloheximide (Sigma-Aldrich) for 4 h. Cells were washed twice with Dulbecco’s Phosphate Buffered Saline (DPBS) (Sigma-Aldrich) and lysed in RIPA buffer (Thermo Fisher Scientific) supplemented with cOmplete EDTA free protease inhibitor (Roche, Basel, Switzerland). Protein concentration was determined using the BCA assay (Thermo Fisher Scientific).

For analysis of cell lysates, SKOV3-cells were cultured in McCoy’s 5A Medium supplemented with 10% FCS (Thermo Fisher Scientific). HROC24-cells were cultured in DMEM/Ham’s F12 (Thermo Fisher Scientific) supplemented with 10% FCS and 3 mM glutamine (Thermo Fisher Scientific). Jurkat subclone JMP cells were cultured in RPMI (Thermo Fisher Scientific) supplemented with 10% NCS (Thermo Fisher Scientific). Huvec cells were cultured in Medium 199 without phenol red (Thermo Fisher Scientific) supplemented with 6.6 % FCS and 33 % EBM™-2 Endothelial Cell Growth Basal Medium-2 (Lonza, Basel, Switzerland). HELA cells (DSMZ ACC-57) and HEK293 cells (DSMZ ACC-305) were cultured in DMEM (Thermo Fisher Scientific) supplemented with 10% FCS. THP-1 cells (ATCC TIB-202) were cultured in RPMI supplemented with 10% FCS. Cells were washed in DPBS (Sigma-Aldrich), before being lysed in M-PER buffer (Thermo Fisher Scientific) supplemented with protease inhibitor cOmplete EDTA-free (Roche) and Phosphatase Stop (Roche). Protein concentration was determined using the Bradford assay (Thermo Fisher Scientific).

### Immunoprecipitation

AMPylated proteins were precipitated from purified recombinant Bip and Bip-AMP or Cho-K1 lysates stressed with either thapsigargin or cycloheximide with antibody 17G6 using Pierce ChIP-grade Protein A/G Magnetic Beads (Thermo Fisher Scientific) according to the manufacturer’s protocol. In short, in a total volume of 500µl 20 µg recombinant protein or 1 mg total protein lysate were incubated with 10 µg 17G6 antibody in 25 mM Tris pH 7.4, 150 mM NaCl, 1 mM EDTA, 5 % glycerol, 1 % NP40 overnight, before being precipitated with 25 µl equilibrated beads. AMPylated proteins were eluted with 100 µl 1x Laemmli for 15 min at 30 °C. Lysate elutions were concentrated to 20µl in a SpeedVac alpha RVC (Christ) without heat. For recombinant protein samples 7.5 µl each of input and unbound sample supplemented with 5x Laemmli buffer and 2.5 µl elution, for lysate samples 5 µl each of input and unbound sample supplemented with 6x Laemmli buffer and 20 µl of concentrated elution were analyzed by 12% SDS PAGE and WB as described.

## Acknowledgements

SKOV3-cells were a kind gift of Leticia Oliveira-Ferrer, Department of Gynecology, University Medical Centre Hamburg-Eppendorf. HROC24-cells were a kind gift of Yuan-Na Lin, Department of General, Visceral and Thoracic Surgery, University Medical Centre Hamburg-Eppendorf. Jurkat cells, subclone JMP (DMSZ verified), and ChoK1-FlpIn cells were a kind gift of Ralf Fliegert, Institute of Biochemistry and Molecular Cell Biology, University Medical Centre Hamburg-Eppendorf. HUVEC cells isolated from umbilical cords were a kind gift of Volker Huck, Centre for Internal Medicine / Diagnostics, University Medical Centre Hamburg-Eppendorf in cooperation with Kurt Hecher, Department of Obstetrics and Fetal Medicine, University Medical Centre Hamburg-Eppendorf. Dorothea Höpfner was supported by a PhD scholarship of the Konrad-Adenauer-Stiftung. We are grateful to the Knut and Alice Wallenberg Foundation, Sweden (KAW 2013.0187 to C.H.), and the Swedish Research Council (VR) for generous support.

## Supplement

### Antibody generation and selection of antibodies binding to native epitopes

After three boosts, the sera of immunized mice were evaluated for their ability to recognize the peptide hapten conjugate and native Cdc42-Thr-AMP in ELISA. Only positive hits were evaluated for their WB performance on various AMPylated proteins to check for backbone independent recognition and potential side chain bias. Of ten immunized mice, five of these were able to recognize native Cdc42-Thr-AMP in ELISA as good as the AMPylated hapten without showing binding of Cdc42 alone. All of them were positive against all tested proteins in WB, although no preference for AMPylation at threonine side chains was observable and all sera reacted with tyrosine as well as threonine modifications.

After the final bleed two positive animals, both C57 BL/6 mice, with superior recognition of native and denatured targets in WB and ELISA and no discernible background against unmodified proteins and peptides were selected to perform cell fusion to hybridoma cells. Evaluation using ELISA resulted in 10 clones that were able to recognize native Cdc42-Thr-AMP and were subsequently chosen for WB testing as described above. The variation of performance among the clones proved to be a lot higher than between mice sera samples, with many clones showing a lack of universal recognition of all targets, high background or strong differences in strength of recognition depending on the AMPylated protein.

Two promising clones with similar very good recognition of all AMPylated proteins in WB independent of their modified side chain, native recognition of Cdc42-Thr-AMP in ELISA and low background were selected for subcloning and subsequent production and purification. One further clone was selected for its unexpected development of a tyrosine-specific recognition, despite immunization with a threonine-modified peptide. Interestingly, antibody 1G11 lost its clear preference for tyrosine AMPylation after upscaling for production (Figure 2A). However, tyrosine-specific recognition of 1G11 could be sharply enhanced in the presence of 1 mM MnCl_2_ (Figure 3B).

In our experience, subcloning and upscaling for antibody production poses the risk of losing binding abilities. One clone lost its AMPylation recognition abilities during subcloning and had to be recloned (Switch in isotypes from IgG1 to IgG2b). Another clone lost its performance during upscaling for antibody production and had to be redone. Therefore, rigorous retesting after each step is crucial.

**Figure S1:**
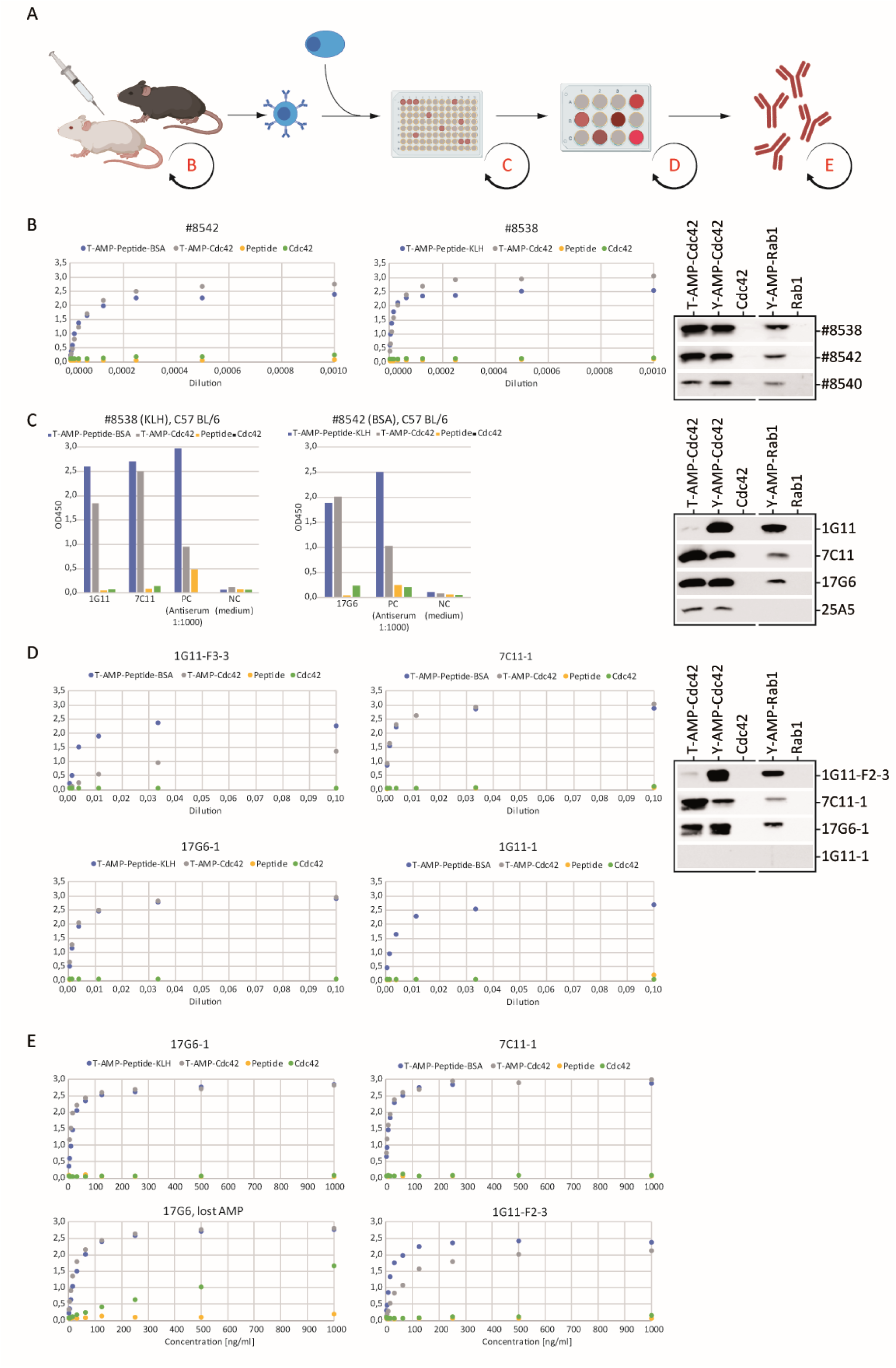
Antibody generation and selection of antibodies binding to native epitopes. **A** Immunization protocol of mice. Evaluation time points are indicated by circular arrows. **B-D** Evaluation regarding recognition of AMPylation in WB and ELISA of **B** mice sera **C** parental hybridoma clones **D** subclones and **E** produced and purified antibodies.

### analytical SEC: supplemental data to Figure 2E

**Figure S 2:**
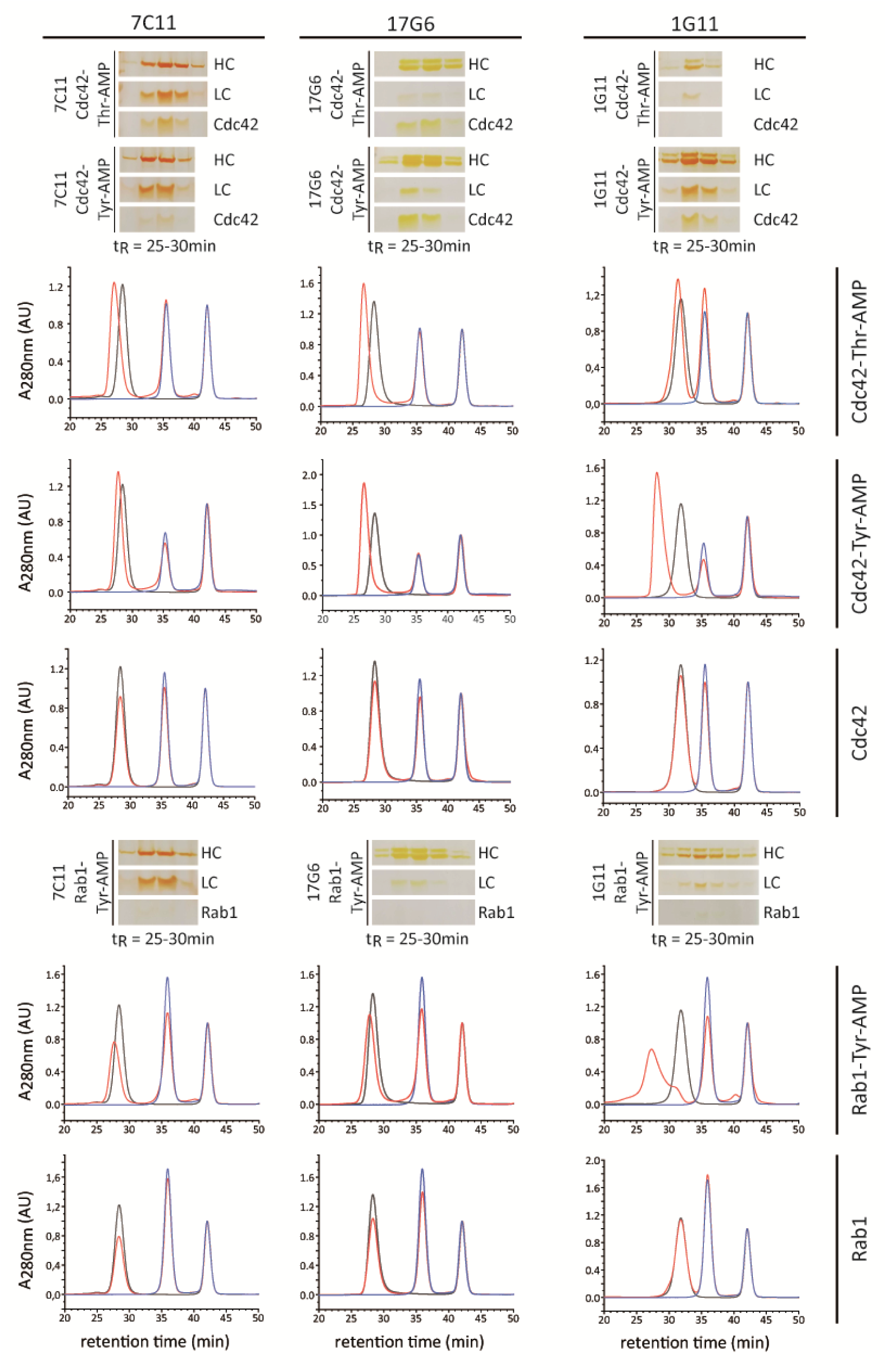
Native binding of AMPylated antigens by all three monoclonal antibodies as determined by analytical size exclusion chromatography. In black antibody alone, in blue antigen alone as indicated, in red co-incubation of antibody and antigen as indicated. Shifted antibody peaks (red) upon co-incubation with AMPylated antigens were fractionated and analyzed by silver stained SDS PAGE.

### MS verification of NMP incorporation

**Figure S3:**
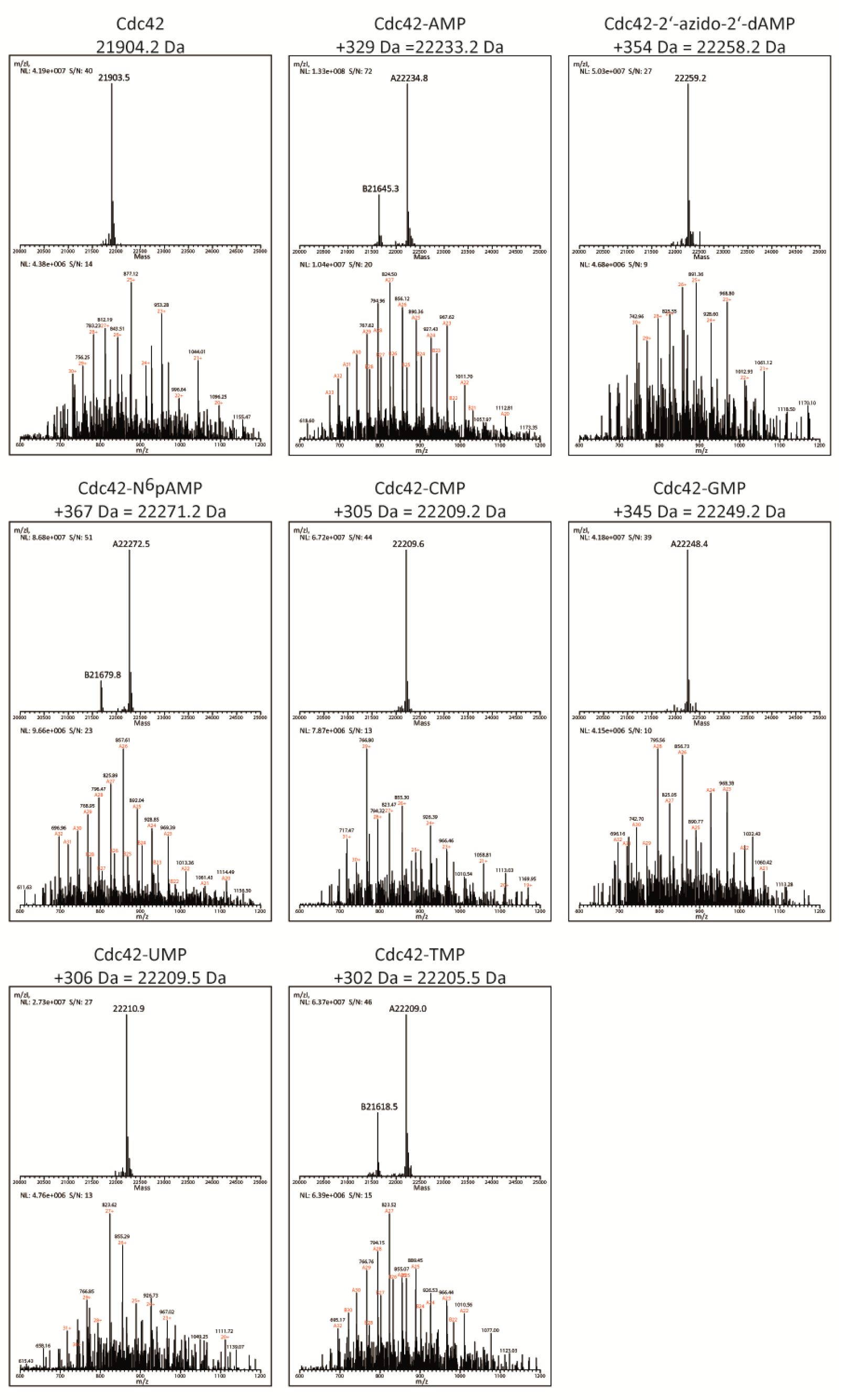
Mass spectrometry confirmation of NMP incorporation by IbpA into Cdc42. Expected molecular weights are indicated above spectra.

### Extended data to cell lysates in Figure 4D

**Figure S4:**
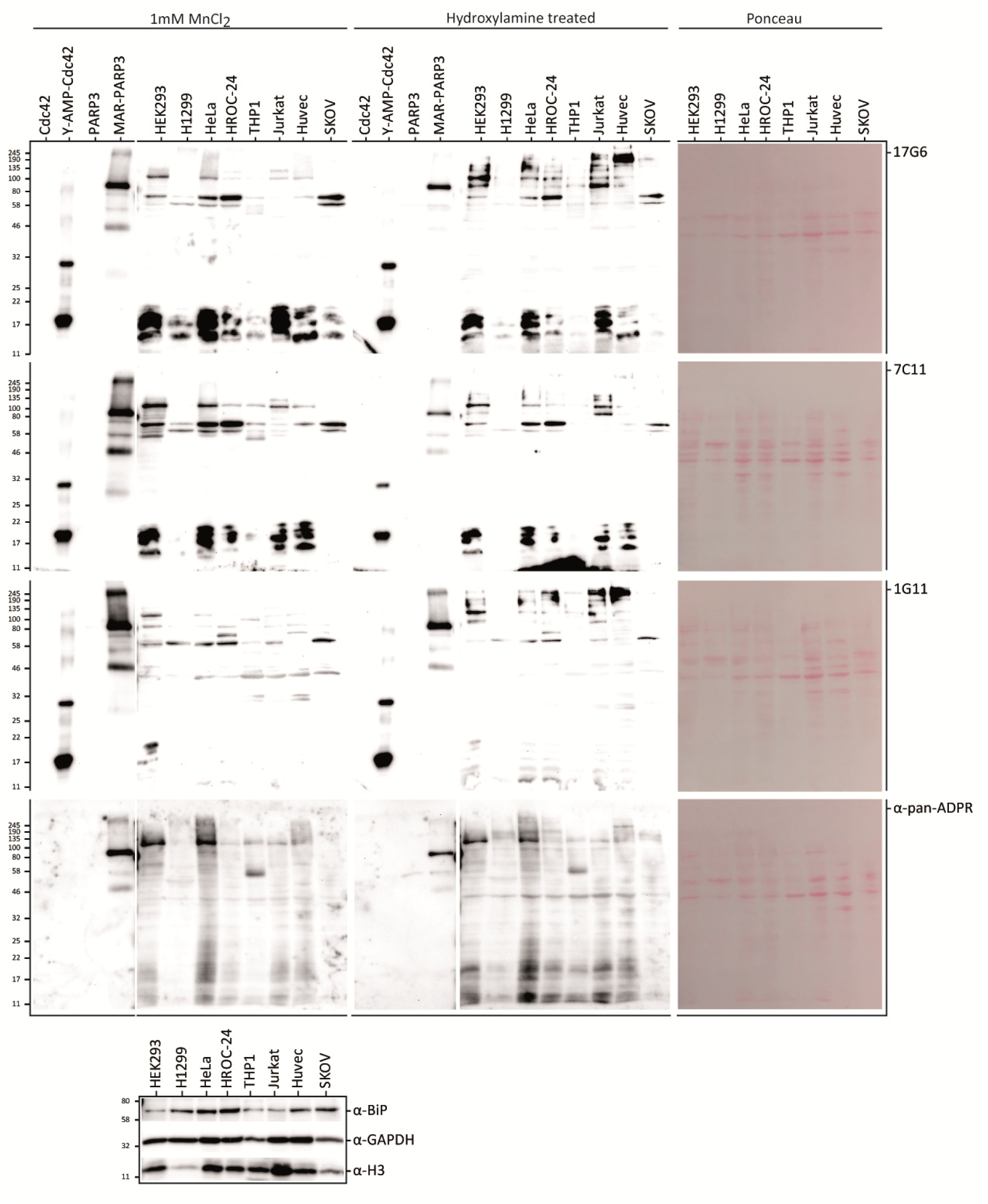
Using all three monoclonal anti-AMP antibodies on various immortalized and cancer cell lines reveals diverse cellular AMPylation. 20 µg cell lysate per lane as indicated were blotted and probed with antibodies as indicated using 1 mM MnCl_2_ as additive. Afterwards cells were treated with 1 M hydroxylamine for 8 h at room temperature to cleave ADP-ribosylation at aspartate and glutamate residues and reprobed with antibody 17G6 using 1 mM MnCl_2_. Antibodies against Bip, GAPDH and Histone H3 serve as loading control. 50 ng recombinant Cdc42-Tyr-AMP serve as positive ctrl for AMPylation, 50 ng recombinant MAR-PARP3 as positive ctrl for ADP-ribosylation and successful hydroxylamine treatment. 50 ng unmodified counterparts are included as negative ctrl.

### Synthesis of Thr-AMP hapten peptide S2 and Thr peptide S3

**Figure S 5:**
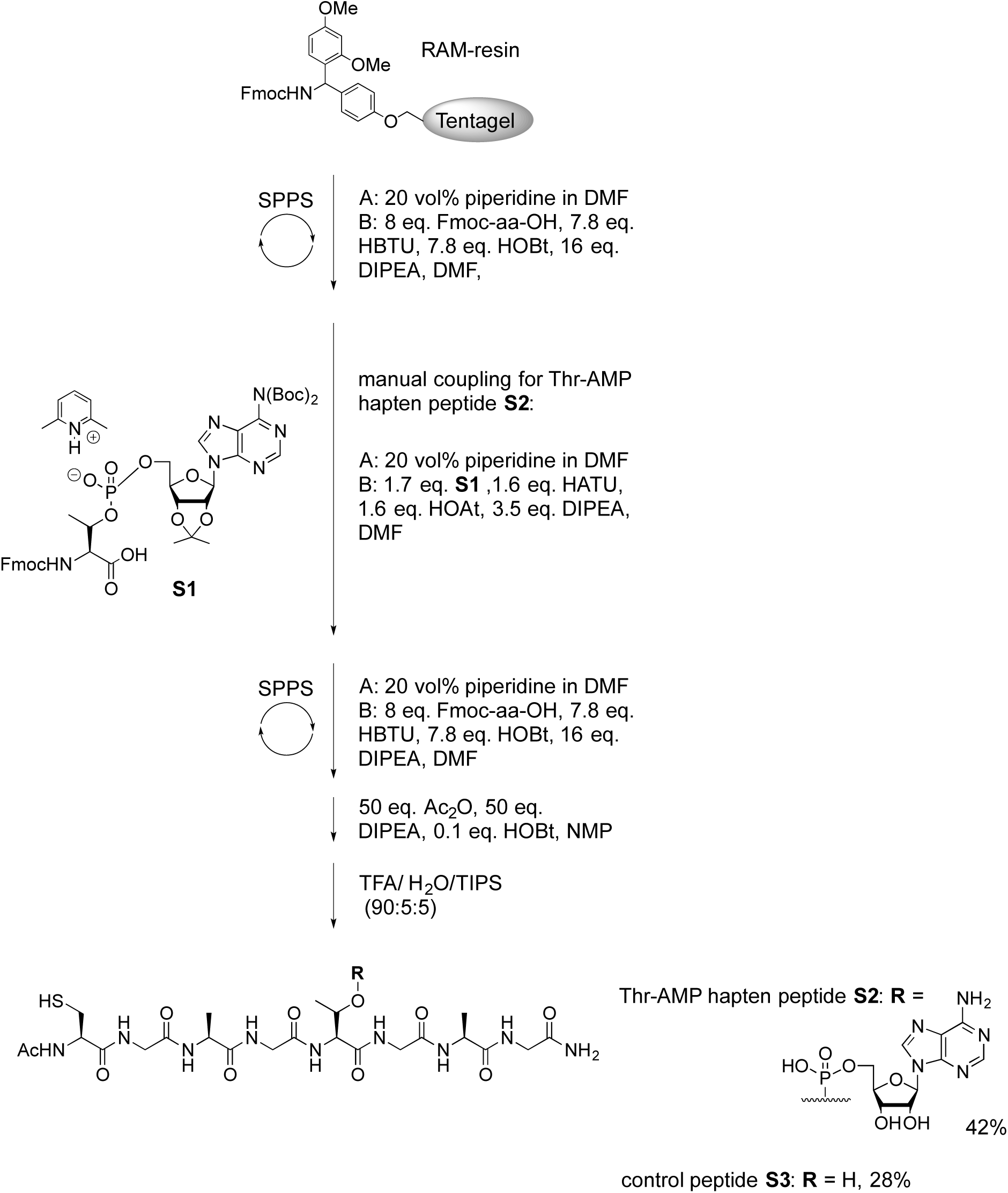
Solid phase peptide synthesis scheme of peptides **S2** and **S3**.

### Thr-AMP hapten peptide Ac-CGAGT(AMP)GAG-NH_2_ (S2)

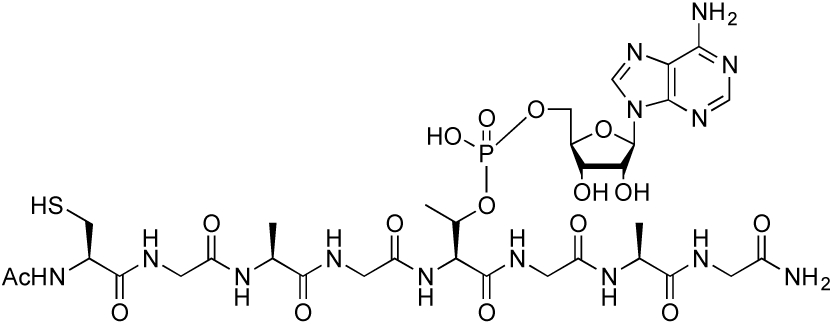

Thr-AMP hapten peptide **S2** was synthesized on 100 mg (25 μmol) of Tentagel-S-RAM resin.

Yield: 42% (10.0 mg, 10.4 μmol).

Analytical HPLC R_t_ = 2.651 min (A/B, (98 : 2) → (75 : 25), 500 μL/min, 7 min); Preparative HPLC R_t_ = 8.945 min (A/B, (97.5 : 2.5) → (50 : 50), 20 mL/min, 10 min).

HR-ESI-MS, m/z: 482.1616 ([M+2H]^2+^, calc. 482.1606), 963.3260 ([M+H]^+^, calc. 963.3139).

^1^H NMR (600 MHz, D_2_O) δ: 8.52 (s, 1H), 8.35 (s, 1H), 4.62-4.59 (m, 1H), 4.43-4.39 (m, 3H), 4.31-4.27 (m, 2H), 4.18 (q, J = 7.3 Hz, 1H), 4.04-3.97 (m, 2H), 3.94 (s_br_, 2H), 3.87 (s, 2H), 3.84 (s, 2H), 3.79 (d, J = 5.2 Hz, 2H), 2.83 (d, J = 6.0 Hz, 2H), 1.98 (s, 3H), 1.31 (d, J = 7.3 Hz, 3H), 1.28 (d, J = 7.3 Hz, 3H), 1.22 (d, J = 6.4 Hz, 3H).

^31^P NMR (242.9 MHz, D_2_O) δ: -1.31 (s).

^13^C NMR (125.9 MHz, D_2_O) δ: 175.50, 175.39, 174.50, 174.13, 172. 79, 171.80, 171.61, 171.21, 170. 95, 150.25, 148.09, 142.24, 118.64, 118.64, 87.97, 84.01, 83.95, 74.49, 72.02, 71.99, 70.19, 64.72, 64.68, 58.43, 58.38, 55.73, 49.94, 49.55, 42.52, 42.45, 42.34, 42.07, 25.16, 21.69, 17.92, 16.68, 16.19.

### Thr peptide Ac-CGAGTGAG-NH_2_ (S3)

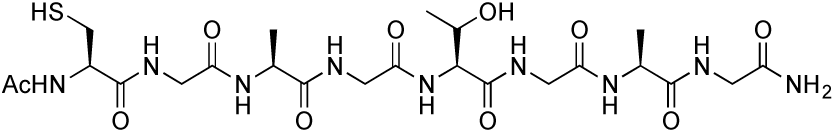

The threonine peptide **S3** was synthesized on 100 mg (25 μmol) of Tentagel-S-RAM resin. Yield: 28% (4.4 mg, 6.9 μmol).

Analytical HPLC R_t_ = 2.449 min (A/B, (98 : 2) → (75 : 25), 500 μL/min, 7 min); Preparative HPLC R_t_ = 7.148 min (A/B, (97.5 : 2.5) → (50 : 50), 20 mL/min, 10 min).

HR-ESI-MS, m/z: 634.2625 ([M+H]^+^, calc. 634.2613), 656.2450 ([M+Na]^+^, calc. 656.2433).

^1^H NMR (600 MHz, D_2_O) δ: 4.43 (t, J = 6.1 Hz, 1H), 4.29-4.18 (m, 4H), 3.94-3.88 (m, 6H), 3.81 (d, J = 5.7 Hz, 2H), 2.85 (d, J = 6.2 Hz, 2H), 1.98 (s, 3H), 1.33-1.30 (m, 6H), 1.12 (d, J = 6.4 Hz, 3H).

^13^C NMR (125.9 MHz, D_2_O) δ: 175.60, 175.51, 174.52, 174.13, 172.81, 172.65, 171.86, 171.36, 171.12, 66.97, 59.14, 58.38, 55.73, 49.92, 49.80, 42.47, 42.45, 42.41, 42.07, 25.15, 21.69, 18.63, 16.47, 16.27.

### NMR spectra of peptides S2 and S3

#### S2 ^1^H-NMR

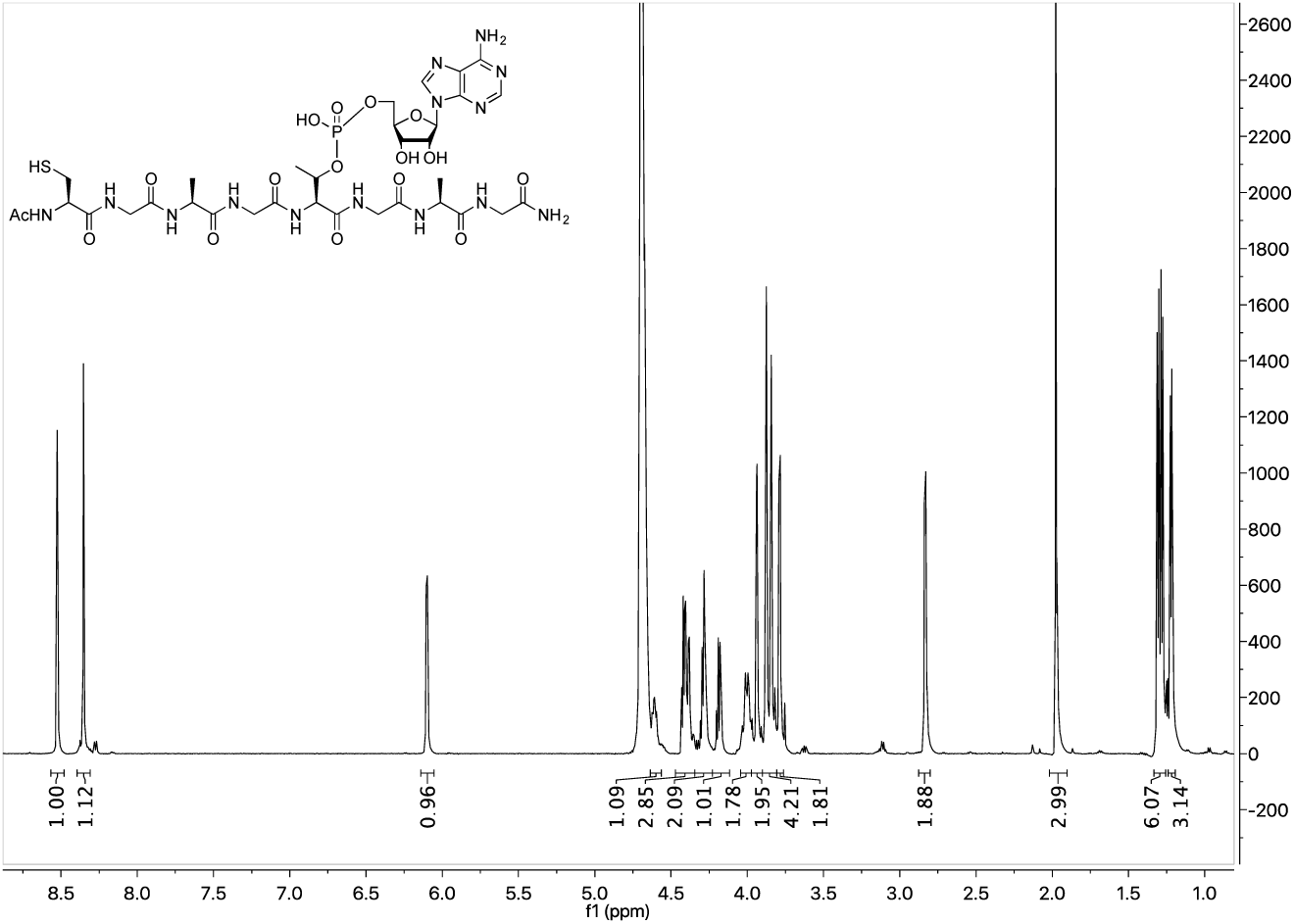

#### S2 ^31^P-NMR

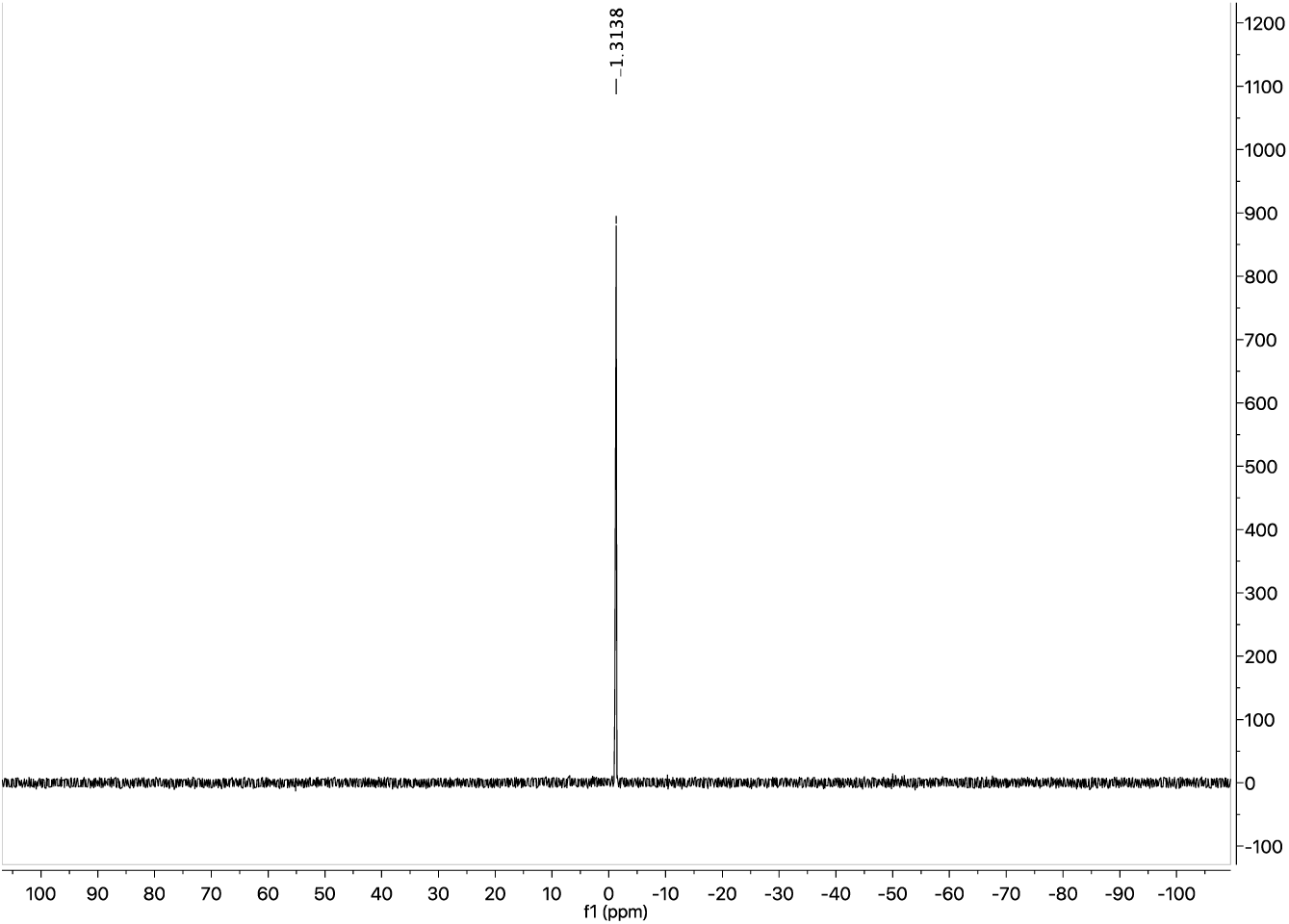

#### S2 ^13^C-NMR

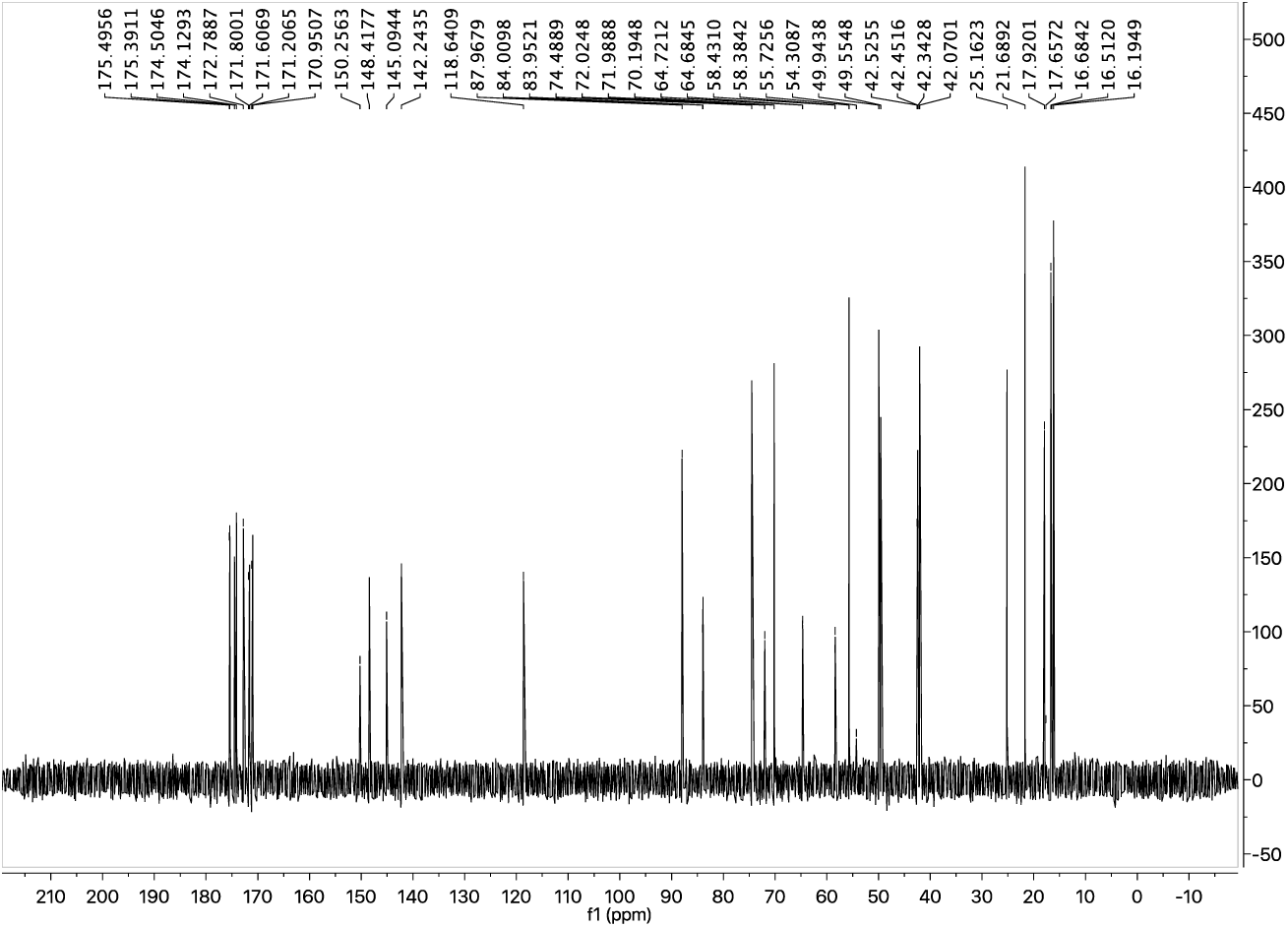

#### S3 ^1^H-NMR

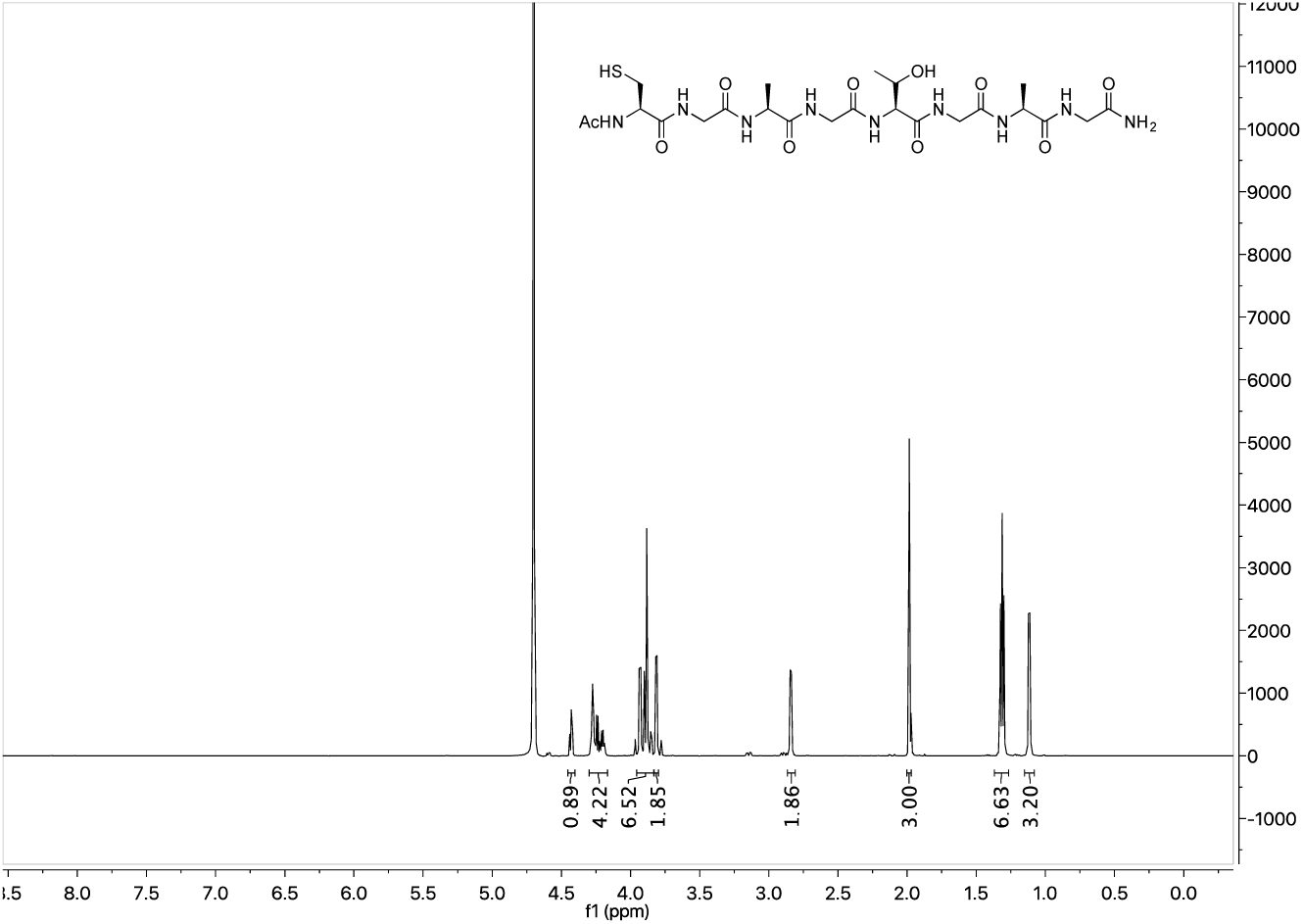

#### S3 ^13^C-NMR

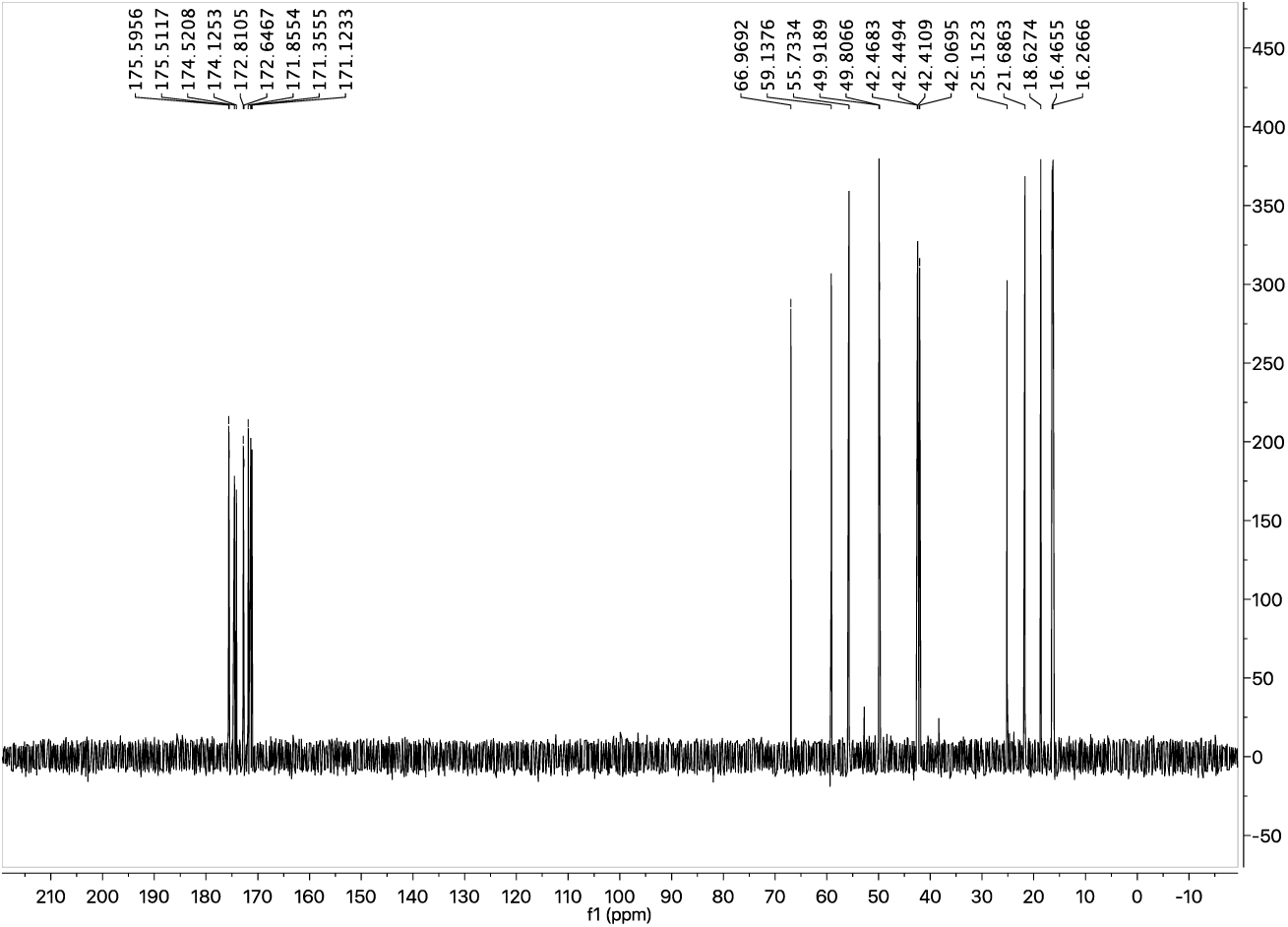

### HPLC chromatograms and HR-ESI-MS spectra of peptides S2 and S3

#### S2 HPLC (UV 214 nm)

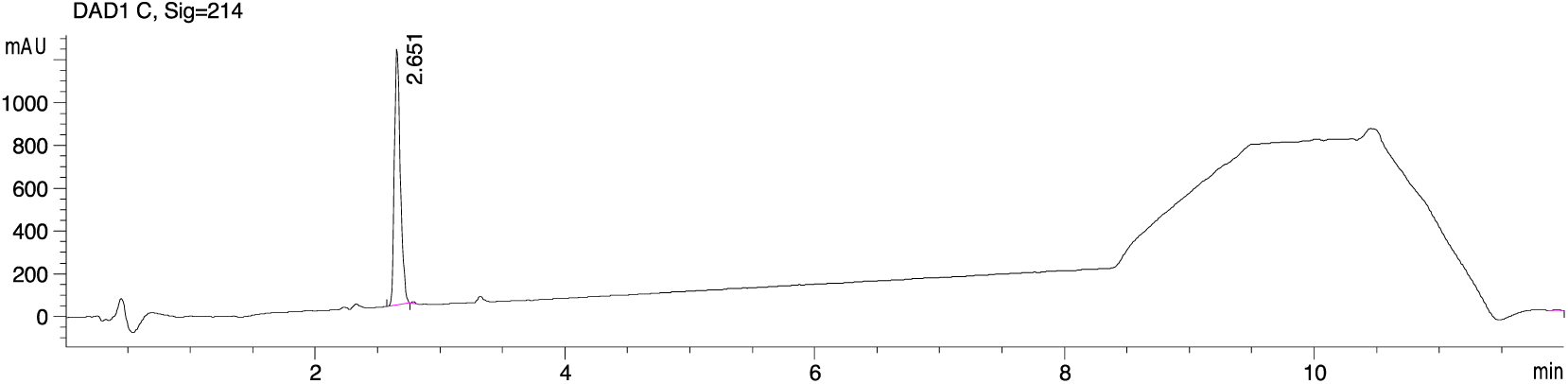

#### S2 HR-ESI-MS

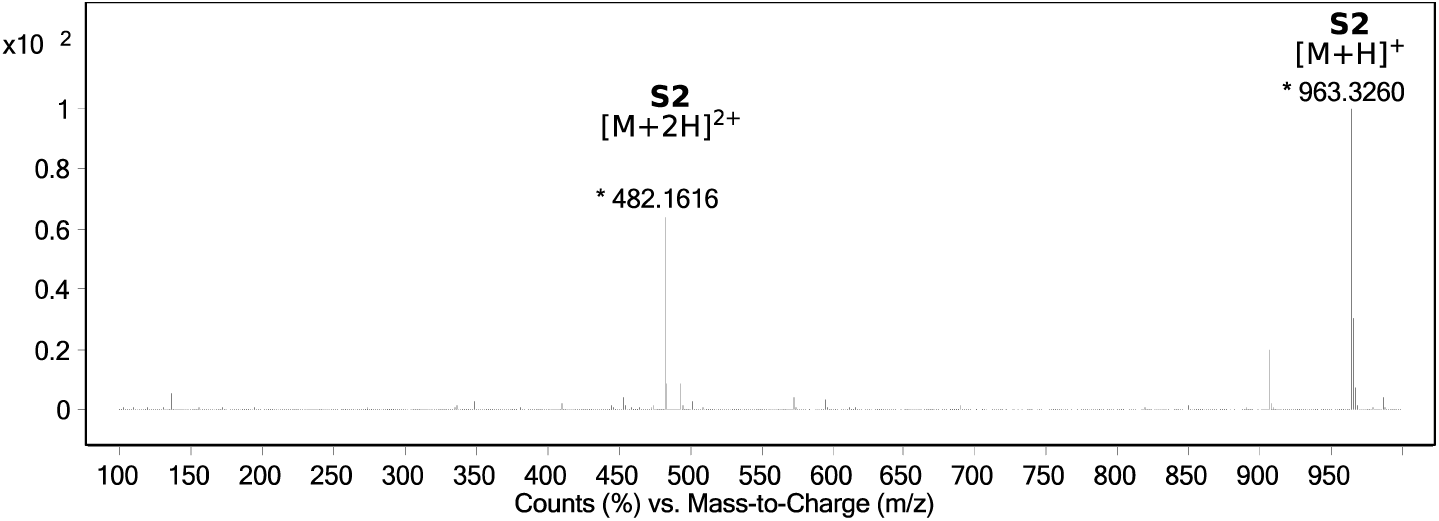

#### S3 HPLC (UV 214 nm)

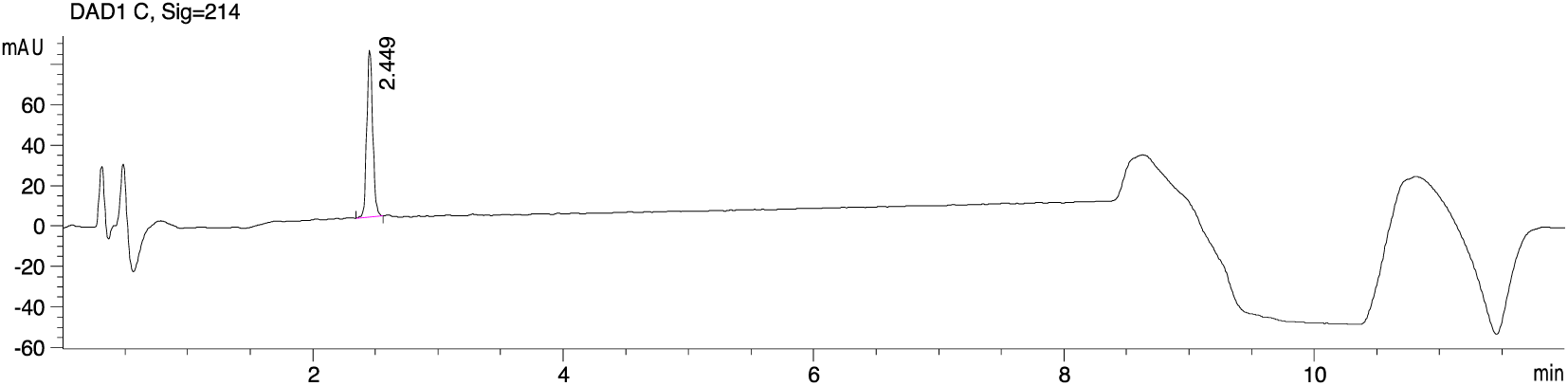

#### S3 HR-ESI-MS

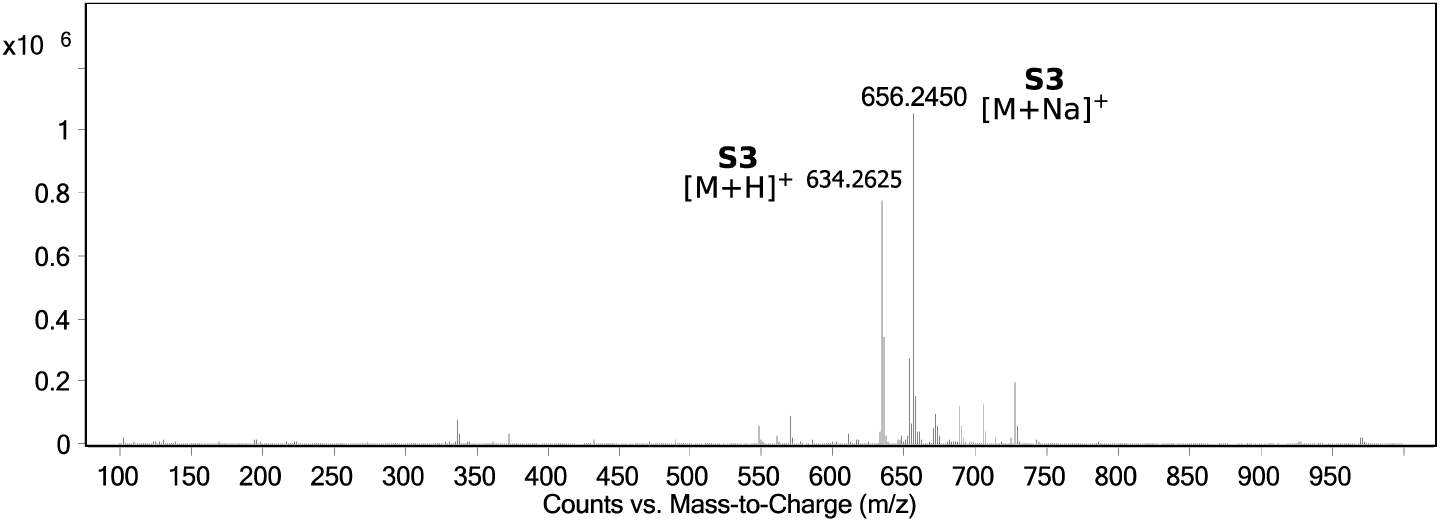

